# Monocytes use protrusive forces to generate migration paths in viscoelastic collagen-based extracellular matrices

**DOI:** 10.1101/2023.06.09.544394

**Authors:** Kolade Adebowale, Cole Allan, Byunghang Ha, Aashrith Saraswathibhatla, Junqin Zhu, Dhiraj Indana, Medeea C. Popescu, Sally Demirdjian, Hunter A. Martinez, Alex Esclamado, Jin Yang, Michael C. Bassik, Christian Franck, Paul L. Bollyky, Ovijit Chaudhuri

## Abstract

Circulating monocytes are recruited to the tumor microenvironment, where they can differentiate into macrophages that mediate tumor progression. To reach the tumor microenvironment, monocytes must first extravasate and migrate through the type-1 collagen rich stromal matrix. The viscoelastic stromal matrix around tumors not only stiffens relative to normal stromal matrix, but often exhibits enhanced viscous characteristics, as indicated by a higher loss tangent or faster stress relaxation rate. Here, we studied how changes in matrix stiffness and viscoelasticity, impact the three-dimensional migration of monocytes through stromal-like matrices. Interpenetrating networks of type-1 collagen and alginate, which enable independent tunability of stiffness and stress relaxation over physiologically relevant ranges, were used as confining matrices for three-dimensional culture of monocytes. Increased stiffness and faster stress relaxation independently enhanced the 3D migration of monocytes. Migrating monocytes have an ellipsoidal or rounded wedge-like morphology, reminiscent of amoeboid migration, with accumulation of actin at the trailing edge. Matrix adhesions were dispensable for monocyte migration in 3D, but migration did require actin polymerization and myosin contractility. Mechanistic studies indicate that actin polymerization at the leading edge generates protrusive forces that open a path for the monocytes to migrate through in the confining viscoelastic matrices. Taken together, our findings implicate matrix stiffness and stress relaxation as key mediators of monocyte migration and reveal how monocytes use pushing forces at the leading edge mediated by actin polymerization to generate migration paths in confining viscoelastic matrices.

**Significance Statement:** Cell migration is essential for numerous biological processes in health and disease, including for immune cell trafficking. Monocyte immune cells migrate through extracellular matrix to the tumor microenvironment where they can play a role in regulating cancer progression. Increased extracellular matrix (ECM) stiffness and viscoelasticity have been implicated in cancer progression, but the impact of these changes in the ECM on monocyte migration remains unknown. Here, we find that increased ECM stiffness and viscoelasticity promote monocyte migration. Interestingly, we reveal a previously undescribed adhesion-independent mode of migration whereby monocytes generate a path to migrate through pushing forces at the leading edge. These findings help elucidate how changes in the tumor microenvironment impact monocyte trafficking and thereby disease progression.

## Introduction

Immune cell migration plays an important role during inflammation and cancer progression(1, 2). Monocytes, a subset of immune cells, are disproportionately recruited from the blood stream during these processes, and they migrate through the extracellular matrix (ECM) to reach the tumor where they can differentiate into macrophages or dendritic cells (3). Cancer progression is often associated with changes in the mechanical properties of the viscoelastic ECM, suggesting that the impact of these changes on monocyte migration could be significant to cancer progression. Specifically, mechanical properties of the ECM such as stiffness, viscoelasticity, and collagen fiber architecture, are altered during malignancy for certain cancers. For example, in breast cancer, stiffness increases from ∼0.1 – 1 kPa in normal tissues to ∼1 – 10 kPa for in malignant tumors(4–7). In addition, some tumors exhibit greater viscous-like behavior compared to normal tissues(4, 8, 9). In biological tissues and ECMs, the viscous resistance, as measured by the loss modulus, is typically around 10% of the elastic resistance, as measured by the storage modulus, at 1 Hz, thus exhibiting a loss tangent of 0.1(10). One consequence of this viscoelastic behavior is that the resistance of the matrix to deformation is reduced over time, a behavior termed stress relaxation. The characteristic timescales of stress relaxation in tissues can range from greater than 1,000 seconds to tens of seconds(10). Lastly, the architecture of the collagen-rich, stromal matrix surrounding tumors changes from wavy to linearized architecture(11). The impact of these changes in matrix mechanics on the three-dimensional migration of monocytes remains unknown.

Cell migration is typically characterized as either amoeboid or mesenchymal based on cell morphological characteristics and activity of cytoskeletal and adhesive machinery(12, 13). Amoeboid migration is characterized by rounded or ellipsoidal cell morphologies, weak cell-ECM adhesions, low proteolytic activity, and Rho-mediated contractility(12). Rho-mediated contractility, acting through Rho kinase (ROCK), is thought to be critical for squeezing the stiff nucleus through confining pores, and prior studies have shown that some immune cells, including T-cells, dendritic cells, and neutrophils are capable of integrin-independent motility(14–18). On the other hand, mesenchymal migration involves more elongated cell morphologies, requires strong adhesions and high proteolytic activity, and can be independent of ROCK activity(12, 19). Furthermore, mesenchymal migration, and to a lesser extent amoeboid migration, can require generation of contractile traction forces on the substrate(20–25). While macrophages and neutrophils exhibit both amoeboid and mesenchymal migration characteristics, the morphologies monocytes utilize to migrate remain unclear(2, 26). Furthermore, much of our current understanding of cell migration is based on cell migration on 2D surfaces, microporous collage gels with sufficiently large pore sizes, or micropatterned confined environments where a migration path is pre-existing(27). In contrast, immune cell migration in vivo occurs in a three-dimensional context that is often confining, and pre-existing migration paths are not always present.

Here, we investigated the impact of changes in matrix stiffness and viscoelasticity on the three-dimensional migration of monocytes, and how monocytes generate paths to migrate in confining matrices. Interpenetrating networks of alginate and type-1 collagen (IPN) with independently tunable stiffness and stress relaxation are developed to model the type-1 collagen rich stromal matrix. Enhanced stiffness and faster stress relaxation both promote the migration of monocytes. Monocytes migrated using a wedge or ellipsoid morphology and depended on actin polymerization and myosin-based contractility. Cells also migrated when encapsulated in non-adhesive viscoelastic matrix consisting of alginate alone with similar characteristics, indicating the dispensability of adhesion for migration under non-adhesive conditions. Mechanistically, cells migrated by generating protrusive forces at the leading edge of the cell, pushing off of the trailing edge or cell sides, which open a migration path in the confining matrix. Together, these results establish matrix viscoelasticity as a key regulator of monocyte cell migration and reveal how monocytes generate paths to migrate in cells in confining matrices.

## Results

### Development of stromal-like matrices with independently tunable stiffness and viscoelasticity

We developed interpenetrating networks of type-1 collagen and unmodified alginate (IPNs) with independently tunable stiffness and viscoelasticity as a model of the stromal matrix(28). The type-1 collagen network mimics the collagen structure found in the type-1 collagen rich stroma, while the alginate network enables tunability of the overall IPN mechanical properties (Fig. 1A, B). Alginate does not provide any adhesion motifs for cells to bind to and is not susceptible to degradation by mammalian proteases(29). IPN stiffness was increased from 1 kPa to ∼2.5 kPa by increasing the amount of calcium crosslinker (Figs. 1C,D). Matrix stress relaxation was modulated from ∼100 s (fast relaxing) to ∼1,000s (slow relaxing), corresponding to a loss tangent of ∼0.12 and ∼0.08 respectively, by utilizing high or low molecular weight alginate though keeping the overall mass density of alginate constant (Figs. 1E,F. Fig. S1). Note that the strategy to modulate viscoelasticity used here is different from a previous approach with collagen-alginate IPNs that used covalent versus ionic crosslinks where the alginate was chemically modified with norbornene and tetrazine groups to facilitate click chemistry(30). The range of stiffness and stress relaxation developed is generally relevant to what is observed during breast cancer, pancreatic cancer, and bladder cancer progression(4, 9, 31). The collagen structure, visualized via confocal reflectance microscopy, showed a fiber architecture with micron-scale spacing between fibers (Fig. 1G). The nanoporous alginate mesh is expected to fill the space between the fibers, making these matrices highly confining. Collagen fiber length and width measurements revealed only some minor differences that are not expected to be biologically meaningful (Figs. 1H-K). Taken together, these data demonstrate the development of collagen-alginate IPNs with tunable stiffness and viscoelastic properties with similar collagen fiber architectures.

**Fig. 1.**
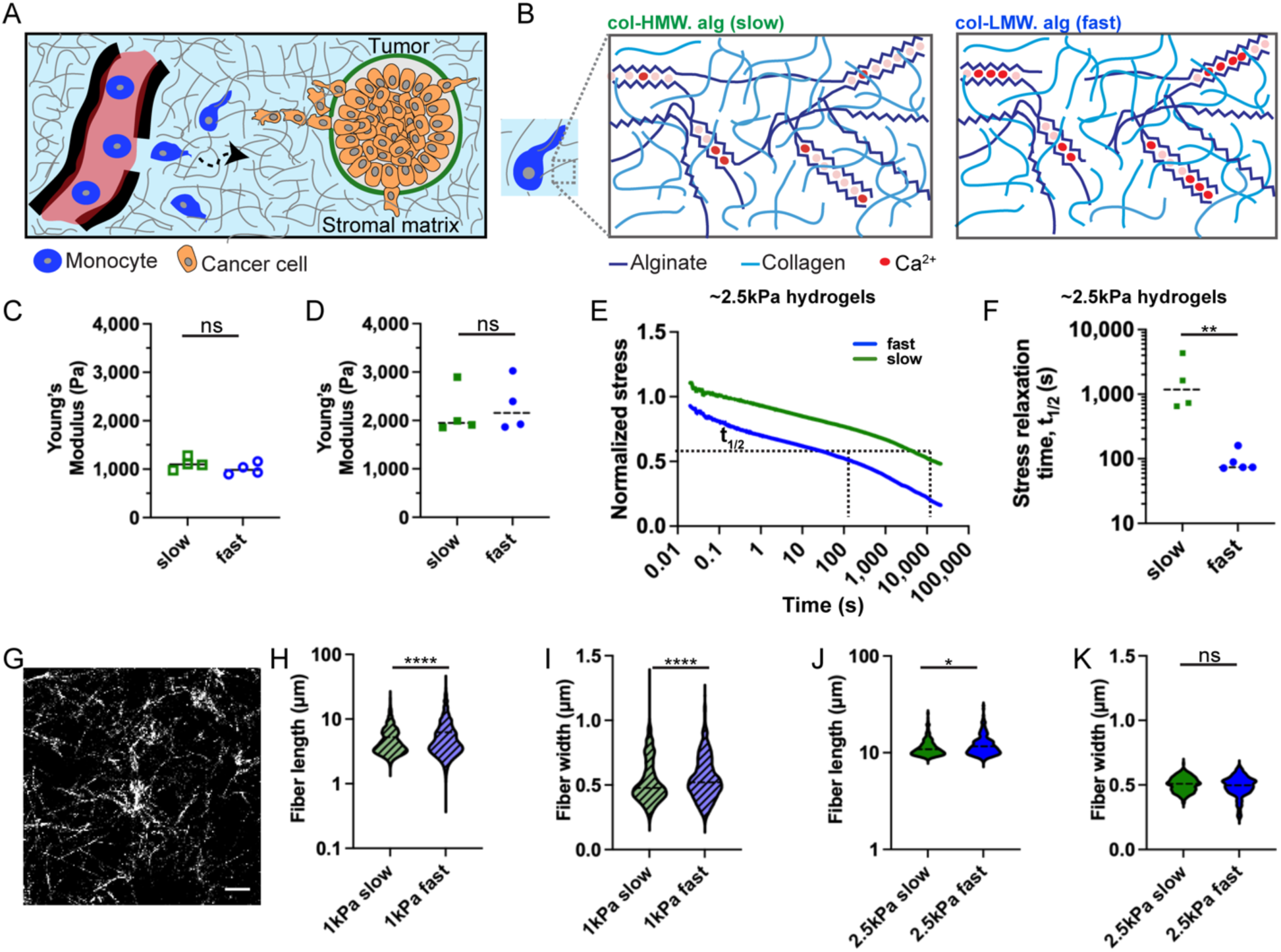
Interpenetrating networks (IPNs) of alginate and collagen with independently tunable properties are used to model the type-1 collagen rich stromal matrix around tumors. (A) Schematic describing how monocytes recruited from the blood migrate through stromal matrix to reach the tumor microenvironment. (B) Schematic of cell encapsulated in IPN made from high molecular weight (HMW) alginate (slow) and low molecular weight (LMW) alginate (fast). (C,D) Young’s modulus measurements of the different IPN formulations d, N = 4 biological replicates for slow and fast. Unpaired t-test; ns p=0.2544, p=0.7130. (E) Representative stress relaxation curves for slow and fast IPNs. (F) Timescale over which the stress relaxes to half its original value for slow and fast relaxing IPNs. N = 4 and 5 for slow and fast respectively. Unpaired t-test; **p=0.0100. (G) Confocal reflectance microscopy of collagen fiber images for IPN. Scale bar: 20 μm. (H-K) Fiber length and width for indicated IPN formulations. N = 2 biological replicates for all conditions. Kolmogorov-Smirnov (due to non-normal distributions); ****p<0.0001, *p=0.0254, ns p = 0.4098. Figs. 1A,B were adapted from previous work(29, 57).

### Increased stiffness and faster stress relaxation promote monocyte migration

With the independently tunable collagen-alginate IPNs, we examined the impact of increased stiffness and faster stress relaxation, independently, on monocyte migration. U937 human monocytes or primary human monocytes were encapsulated in 3D matrices and tracked during time-lapse microscopy experiments (Fig. 2A). Monocytes were generally rounded, but more elongated in the fast relaxing IPNs compared to slow relaxing IPNs (Fig. 2B, Fig. S2). Cell migration speed, mean squared displacement (MSD), and ratio of cell displacement to cell track length (directionality) were used as quantitative descriptors of migration. Increased stress relaxation or stiffness induced longer cell tracks (Figs. 2C-F). Faster matrix stress relaxation increases cell migration speed for the same stiffness values for U937 cells and primary cells from human patients (Figs. 2G-O). Cells also migrated with greater speed in stiffer matrices for both slow and fast relaxing IPNs (Fig. 2J-L). Both the slope and y-intercept of the MSD curve increased with stiffness, indicating that cells migrate more efficiently and with higher free diffusivity as IPN stress relaxation became faster (Fig. 2I). Similar observations were made with increase in matrix stiffness leading to more efficient migrating in fast relaxing IPN (Fig. 2L). Similarly, faster stress relaxation enhanced migration and diffusive behavior of primary monocytes (Figs. 2M-O). Taken together, these results demonstrate that faster matrix stress relaxation and increased stiffness both promote increased migration in monocytes.

**Fig. 2.**
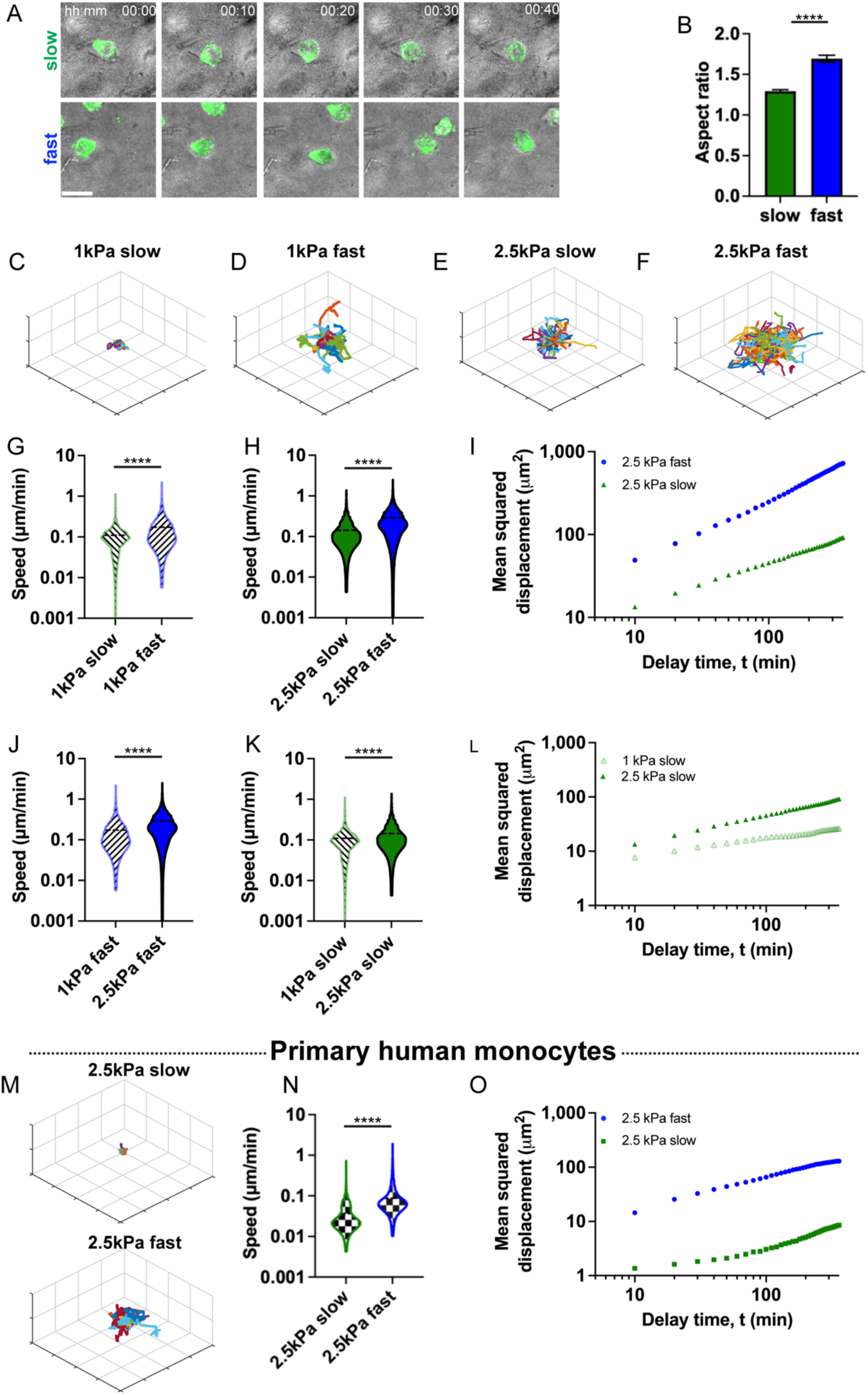
Faster stress relaxation and increased stiffness independently promote enhanced three-dimensional migration of individual monocytes. (A) Representative images of individual U937 cells migrating in slow relaxing (top row) and fast relaxing (bottom row) hydrogels. Scale bar: 20 μm. (B) Aspect ratio of U937 monocytes embedded in slow or fast relaxing matrix. n = 132 cells and N = 2 biological replicates. (C-F) Representative U937 migration tracks for each of the indicated matrix parameters. Each grid is 50 μm. n > 85 for each condition. (G-I), U937 migration speed, and MSD when stress relaxation is enhanced in 1 kPa or 2.5 kPa gels. (G,H), n > 1,148 for each condition; N = 2 biological replicates. (J-L), U937 migration speed, and MSD when stiffness (Young’s modulus) is increased in slow or fast relaxing gels. (J,K) n > 2,288 for each condition; N = 2 biological replicates. (M-O) Representative migration tracks, speeds, and MSD for primary human monocytes in each of the conditions. Each grid is 50 μm. n > 91 for each condition; N = 2 biological replicates. (I,L,O) n > 91; N = 2 biological replicates. (B,G,H,J,K,N) Kolmogorov-Smirnov: **** p < 0.0001.

### Migrating monocytes are amoeboid-like, migrate independent of matrix adhesions

We examined monocyte morphology and the potential role of matrix adhesions to determine the mode of migration. Visualization of cells encapsulated in slow and fast relaxing IPNs revealed that the cells were generally rounded or ellipsoidal, suggesting an amoeboid type of migration (Fig 2A). There was dependence on matrix stress relaxation as the ellipsoidal shape of cells in fast relaxing IPNs was marked by a higher aspect ratio and lower circularity compared to cells in slow relaxing IPNs (Figs. 2B, S2). We wondered if the shape of migrating cells is consistent with known actin structures implicated in migration such as invadopodia and filopodia. Invadopodia are thin actin-rich protrusions typically found at the front of the migrating cell and have an extension-retraction behavior. Filopodia are also thin actin-rich structures that are found protruding from the cell membrane, though not necessarily always associated with migration. We did not observe these invasive protrusions or blebbing morphologies, which are characteristic of certain kinds of mesenchymal and amoeboid migration mechanisms(32).

Cell-matrix adhesions are thought to be important for migration of many cells and critical for mesenchymal migration, though some immune cells are known to be capable of migration independent of adhesions(27). Pharmacological inhibition and CRISPR knockouts (KOs) were applied to perturb the activity of adhesion receptors and associated proteins. Small molecule inhibition of the receptor tyrosine kinase activated in response to collagen, discoidin domain receptor 1 (DDR1), only had a modest impact on cell migration or MSD (Fig. 3A-C). Furthermore, it is thought that monocytes recruited to solid tissues polarize to macrophages whose migration is dependent on heterodimeric α_M_β_2_-integrins(2, 33). Interestingly, CRISPR knockout of β_2_-integrins resulted in a slight increase in cell migration, consistent with what has been observed for human neutrophil migration under confinement(16). But CRISPR knockout of talin-1, an integrin associated protein, did lead to a decrease in cell migration (Figs. 3A-C). CRISPR KO of β_2_-integrins combined with antibody blocking of β_1_-integrins resulted in a decrease in cell migration implicating β_1_-integrins when it comes to monocyte motility in 3D (Fig. 3A, Fig. S3). Differentiation of the monocytes towards a highly adhesive macrophage phenotype by exposure to phorbol myristate acetate (PMA) led to substantial decrease in migration in the collagen-alginate, consistent with the known adhesion-dependent migration of macrophages (2) (Fig. S4).

**Fig. 3.**
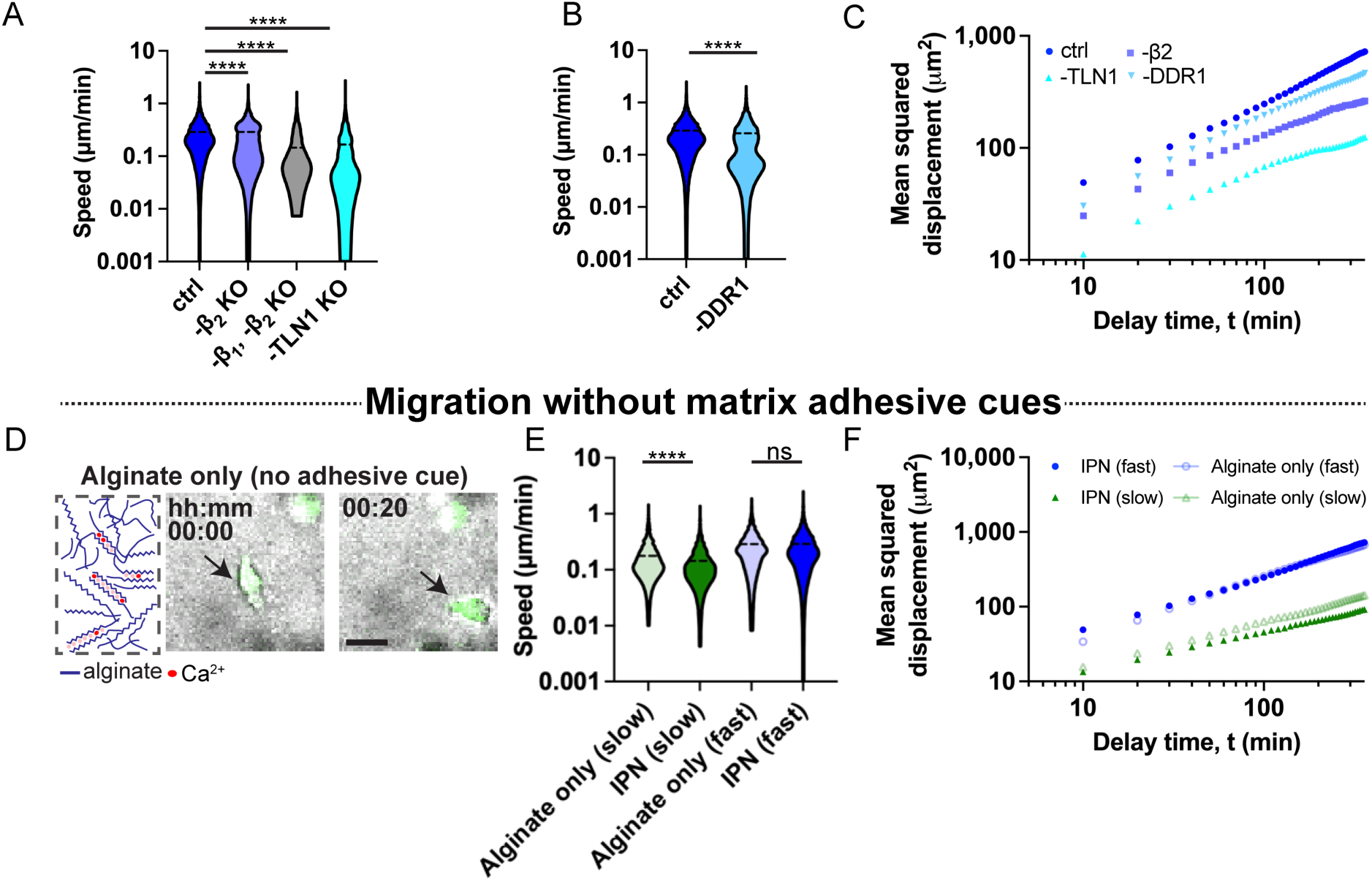
Monocytes migrate in the collagen-alginate IPNs with amoeboid-like morphologies in an adhesion-dispensable manner. (A,B,C) CRISPR/Cas9 mediated knockout of talin-1 decreased migration but inhibition of DDR1 or β2-integrin knockout led to no change or a slight increase in migration speed and MSD. (D) Schematic of alginate matrix without collagen adhesive cue. Typical morphology of a cell migrating in alginate viscoelastic matrix (no collagen) indicated by black arrows. Scale bar: 20 μm. (E) Cell migration speed increases for slow relaxing viscoelastic matrix without collagen compared to the IPN matrix with collagen. Migration speeds are similar for fast relaxing matrix with or without collagen adhesions. (F) Cells display similar MSD profile with or without collagen, indicating similar diffusive behavior. (A) For ctrl, -β2 KO, and -TLN1 n > 2,102; N > 2 biological replicates. For β1,-β2 KO n > 736; N = 2 biological replicates. Kruskal-Wallis test with Dunn’s multiple comparisons: * p = 0.0151, **** p < 0.0001. (B) n > 2,287 for each condition; N > 2 biological replicates. (C) n > 60 for each condition; N = 2 biological replicates. (E) n > 1,606 for each condition; N > 2 biological replicates. (F) n > 145 for each condition; N = 2 biological replicates. (B,E), Kolmogorov-Smirnov: ns p = 0.1096, **** p < 0.0001. Data in this figure are from U937 monocytes.

Given the independence of migration on b2 integrin and DDR1, we sought to further investigate the role of adhesions in monocyte migration. To this end, we performed cell migration studies in pure alginate hydrogels that did not contain any cell-adhesion ligands (Fig. 3D). Monocytes encapsulated in the collagen free viscoelastic matrices had similar morphologies to those in the collagen-1 rich IPN matrices (Fig. 3D). Cells migrated robustly in the pure alginate gels, with a slight increase in speed for monocytes in slow relaxing alginate gels without collagen (Fig. 3E). Furthermore, the MSD showed similar diffusive behavior for cells encapsulated in slow relaxing viscoelastic matrices with or without collagen (Fig. 3F). But there was an increase in MSD diffusive behavior and directionality for cells in fast relaxing matrix without collagen (Fig. 3F). Together, these results confirm cell migration in monocytes to be amoeboid and show that adhesion to the matrix is dispensable for migration.

### Monocyte migration requires actin polymerization, myosin contractility, and WASp

We then investigated the role of actin polymerization, myosin-mediated contractility, and the Rho pathway in mediating monocyte migration, given the known importance of the cytoskeletal machinery in driving cell migration. Actin puncta were observed at the rear of cells (Fig. 4A). The puncta remained localized at the rear of migrating cells while actin was uniformly distributed around the cortex of non-migrating cells (Fig. 4A). Furthermore, immunofluorescence staining revealed activated myosin staining to be somewhat diffuse throughout the cell, indicating that cell-contractility throughout the cell was not localized to any specific region (Fig. 4B). Thus, strong colocalization of actin and myosin was not observed as has previously been described for amoeboid migration(15, 34). Next, actin polymerization, myosin activity, and Rho-mediated contractility were perturbed using pharmacological inhibition to determine their respective roles in cell migration and directionality of migration (Figs. 4C,D). Inhibition of actin polymerization or the Arp2/3 complex, known to drive dendritic actin network growth, led to reduced migration speed, indicating that dendritic actin network growth at the leading edge is critical for cell migration. Similarly, inhibition of Rho-mediated contractility and myosin light chain kinase (MLCK), an activator of myosin, diminished cell migration speed, indicating the role of actomyosin contractility in driving cell migration, consistent with results from previous studies of amoeboid migration(15, 35). We confirmed that pharmacological inhibition did not have a significant impact on cell viability (Fig. S5). Similar results for the impact of pharmacological inhibition on cell migration were obtained for primary monocytes (Fig. S6). In addition, residual primary migration after treatment with latrunculin A was further abrogated by pharmacological inhibition of Na(+)/H(+) exchanger (NHE) ethyl-isopropyl amiloride (EIPA) (Fig. S6), a component that has been recently implicated in cell migration (36).

**Fig. 4.**
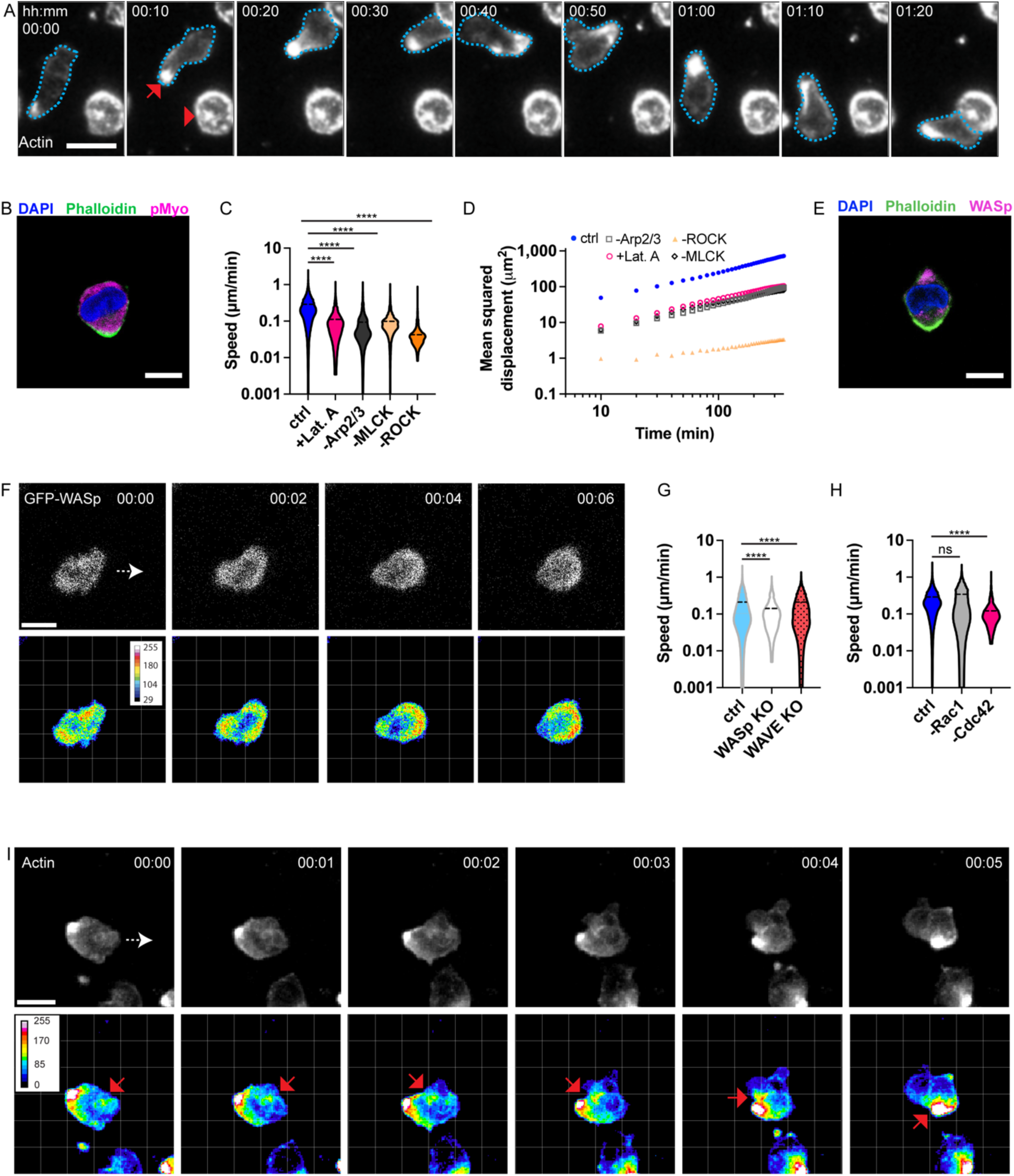
Monocyte migration is dependent on actin polymerization. (A) Actin is localized at rear of migrating cell (red arrow) but uniformly distributed around cell perimeter of a non-migrating cell (arrowhead). Scale bar: 20 μm. (C) Immunofluorescence image of representative cell showing DAPI and phosphorylated myosin II. Dendritic actin nucleator (WASp) and actin filaments (phalloidin) are localized at cell front and rear respectively. Scale bar: 10 μm. (C) Inhibition of actin polymerization, (Lat. A, Arp2/3), myosin activation (MLCK), and Rho contractility (ROCK) decreased cell migration speed. (D) Mean squared displacement (diffusive behavior) of cells under different pharmacological treatment conditions. (E) Immunofluorescence image of representative cell showing DAPI, phalloidin, and WASp. (F) Live imaging of fluorescently labeled WASp at the front of the cell. White arrow indicates migration direction. (G) Effect of CRISPR knockout (KO) of WASp protein and WAVE complex on migration. (H) Effect of Rac1 and Cdc42 RhoGTPases on migration. (I) Live imaging of fluorescently labeled actin. Cell is moving from left to right (white arrow). Red arrow indicates actin can be visualized flowing from the front (right side) to the back of the cell (left side). (C) n > 984 for each condition; N > 2 biological replicates. Kruskal-Wallis test with Dunn’s multiple comparisons: ns p = 0.1496, **** p < 0.0001. (D) n > 139 for each condition; N = 2 biological replicates. (G) n > 1,371 for each condition; N = 2 biological replicates. Kruskal-Wallis test with Dunn’s multiple comparisons: **** p < 0.0001. (H) n > 2,247 for each condition; N = 2 biological replicates. Kruskal-Wallis test with Dunn’s multiple comparisons: ns p > 0.9999, **** p < 0.0001. (E,F,I) Scale bars: 10 μm. Data in this figure are from U937 monocytes.

Given that actin polymerization via the Arp2/3 complex mediates cell migration, we next examined the role of WASp and WAVE, actin nucleation promotion factors that activate the Arp2/3 complex. It has previously been shown that Cdc42 activation leads to activation of WASp, and formation of small protrusions, whereas WAVE activation via Rac1 leads to formation of a growing sheet like protrusion (37–39). Immunohistochemical staining indicates WASp to be localized at the leading edge of the cell, being opposite to the actin puncta at the rear of the cell (Fig. 4E, Figs. S7). To further examine the role of WASp, GFP-WASp was expressed in the cells and imaged during cell migration. WASp was found to be at elevated levels at the leading edge of the cell (Figs. 4F, S8). Further, WASp KO led to a substantial decrease in migration speed (Figs. 4G, S9). In contrast, WAVE KO led to only a marginal decrease in cell migration speed, indicating that it might not play such an important role in migration (Fig. 4G). We then examined the role of Rac1 and Cdc42 RhoGTPases on mediating actin polymerization and cell migration. Cdc42 inhibition strongly inhibited monocyte migration speed, whereas Rac1 inhibition had no impact (Fig. 4H).

While WASp, and presumably actin polymerization, was localized to the leading edge, surprisingly, the strongest actin signal was observed in the dense puncta at the rear of the migrating cell. We reasoned that these two observations could be reconciled if actin polymerization at the leading edge was followed by rearward flow of the actin, leading to accumulation of actin at the rear. Consistent with this expectation, timelapse imaging of actin indicated rearward movement of actin spots relative to the leading edge (Fig. 4I, S10). Together, these data suggest that actin polymerization mediated by the Cdc42-WASp signaling axis drives actin polymerization at the leading edge and cell migration overall (Fig. 4I).

### Cell volume changes are not critical for monocyte migration

Next, we assessed the potential role of cell volume changes on cell migration. Recent studies have implicated volume changes as a potent factor driving migration in some contexts (40–43). Volume was measured before and during migration (Fig. 5A, B). Interestingly, there was an increase in cell volume during cell migration. To assess whether this cell volume increase could mediate migration, cell volume expansion was inhibited through inhibition of TRPV4 mechanosensitive ion channels and Na^+^/H^+^ pumps, using GSK205 and EIPA respectively, due to their previously described roles in volume changes and cell migration. Inhibition of either component abrogated the increase in volume associated with migration (Figs. 5A, B). Despite the lack of volume change with EIPA, cell migration was only very marginally decreased relative to the control, indicating that cell migration and migration path generation can occur robustly without changes in cell volume (Fig. 5C). There was a more substantial decrease in migration speed with inhibition of TRPV4 (Fig. 5D). However, inhibition of TRPV4 decreases stimulated Ca^2+^ influx and could have a pleiotropic effect due to several Ca^2+^-dependent pathways in cells(44). Thus, based on the EIPA data, we conclude that changes in cell volume alone are not critical for driving 3D monocyte migration in our studies.

**Fig. 5.**
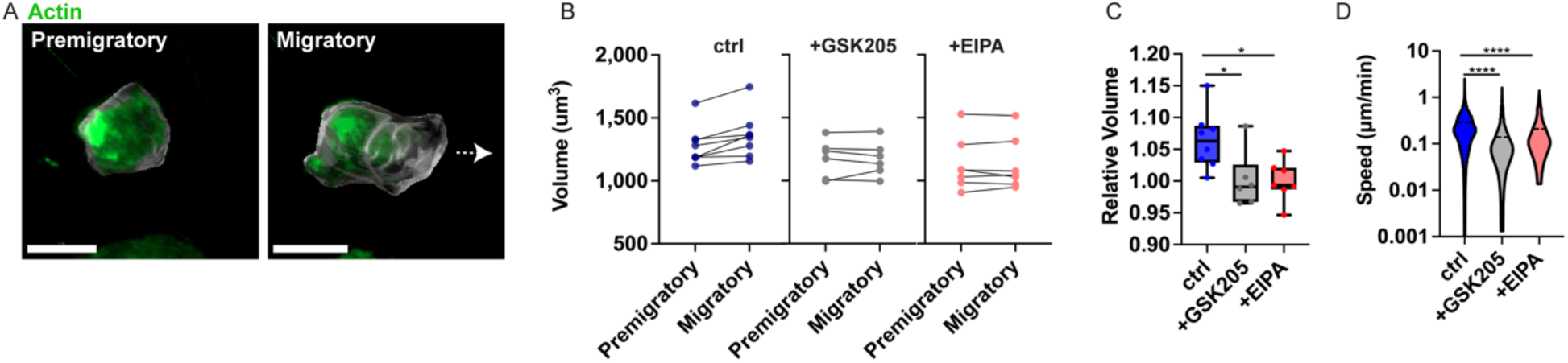
Monocyte migration is not dependent on cell volume changes. (A) Rendered representative cell before and during migration. (B,C) Both EIPA and GSK205 inhibitors negated this volume change. n > 5 for each condition; N = 2 biological replicates. Kruskal-Wallis test with Dunn’s multiple comparisons: * p < 0.05. (D) Both EIPA and GSK205 inhibition led to a decrease in cell migration. n > 1,481 for each condition; N = 2 biological replicates. Kruskal-Wallis test with Dunn’s multiple comparisons: **** p < 0.0001. Scale bar: 10 μm. Data in this figure are from U937 monocytes.

### Actin polymerization generates protrusive forces to drive monocyte migration

The distinct localization of actin at the cell rear and front, and absence of strong colocalization of actin with phosphorylated myosin suggested that cells might utilize actin polymerization to generate pushing forces that open up migration paths in the confining matrices. To investigate this, we analyzed matrix deformation. The three-dimensional ECM deformation field around a single cell showed protrusive deformations at the leading-edge during migration (Fig. 6A). To reduce the computational requirements, ECM deformation in a single optical plane of migrating monocytes was analyzed and an average map of ECM deformations associated with cell migration was generated with an overlay of an average cell shape (Figs. 6B-D(i)). These analyses confirmed the 3D observation of cells applying pushing forces on the matrix at the leading edge, with cells pushing off of their rear and, to some extent, their sides to balance forces. The deformations were abrogated upon inhibition of actin polymerization, myosin contractility, or the Rho pathway (Fig. 6D). Further, migrating cells were observed to generate micron-sized channels in their wake, highlighting that the observed mode of migration generates migration paths in the confining matrices (Fig. 6E).

**Fig. 6.**
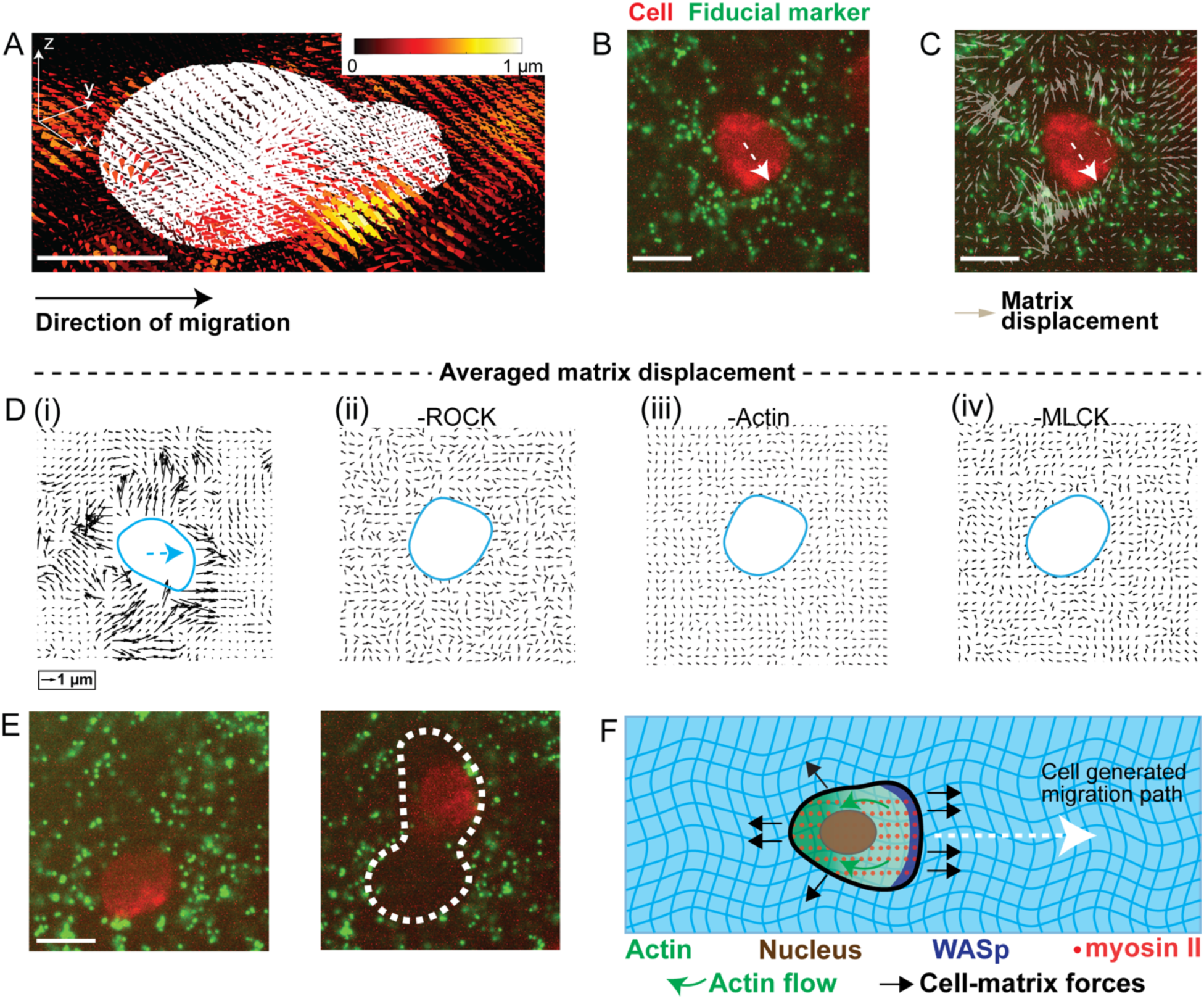
Monocytes apply protrusive forces to generate migration paths in confining IPN matrices. (A) Three-dimensional deformation field around a monocyte. Direction of cones represent the direction of the deformation field. Cell is migrating from left to right. Scale bar: 5 μm. (B,C) Confocal images of fluorescent beads (green) and a monocyte (red). Grey arrows represent ECM deformations generated during migration. White arrow (dotted): direction of migration. (D) Deformation field of ECM induced by migrating monocytes, averaged over 14 cells for (i) and at least 5 cells for (ii)-(iv). N > 2 biological replicates. Cyan arrow indicates monocyte direction of migration, cyan outline indicates cell outline, and black arrows indicate deformation field. (E) Cells open micron-sized channels to migrate. Fluorescent beads (green), and a monocyte (red) are shown 90 min apart. Dotted line (white) indicates the channel created by the monocyte. (F) Proposed model of migration. WASp mediated polymerization generates force to push matrix at the cell front, opening up a channel for migration as cell migrates from left to right. Green arrow: retrograde flow of actin. (B,C,E,F) Scale bar: 10 μm. (D) 2 h (left) and 3.5 h (right) after encapsulation of monocytes in IPNs. Data in this figure are from U937 monocytes.

Taken together, these findings suggest a new mode of migration in monocytes (Fig. 6F). In this mode, actin polymerization generates protrusive forces at the leading edge, with the cells pushing off their trailing edge or sides. The protrusive forces generate micron-size channels in otherwise physically confining matrix for the monocytes to migrate through. This channel generation could occur via divergence of stresses at the leading edge, or via some kind of compression failure in the matrix. Myosin contractility drives flow of the newly polymerized actin at the leading edge towards the rear of the cell, as the cell moves forward.

## Discussion

We studied three-dimensional migration of monocytes in viscoelastic collagen-alginate IPNs. Increased stiffness and faster stress relaxation independently enhance monocyte migration. Monocytes migrate using amoeboid-like morphologies and depend on myosin contractility. However, myosin was diffuse throughout the cell and strong colocalization of myosin with actin was not observed as has been previously described for amoeboid migration(15). Actin polymerization at the leading edge generates pushing forces on the matrix, which opens a migration path, while myosin contractility acts throughout the cell. Together, these results describe a previously undescribed mode of monocyte migration whereby monocytes generate actin mediated protrusive forces at the leading edge to open a path to migrate in confining matrices in 3D.

We developed collagen-alginate IPNs to model the viscoelastic type-1 collagen rich stroma, and capture changes in mechanics associated with cancer progression. Prior work suggest that tissues become stiffer under some pathological conditions. However, recent findings demonstrated that alterations in tissue viscoelasticity are a feature of aberrant tissues(4, 9). Thus, the collagen-alginate IPNs were developed, in which stiffness and stress relaxation (viscoelasticity) were independently tunable, while maintaining a similar type-1 collagen rich architecture. We note that previous studies developed tunable type-1 stromal-matrix mimics using a macromolecular crowding agent (polyethylene glycol) to modulate the network architecture and mechanics of type-1 collagen gels, or click chemistry to covalently crosslink the alginate network in collagen-alginate IPNs, and thereby reduce the stress relaxation rate(30, 45). The approach described here is distinct as neither a crowding agent nor covalent bonds are involved, with IPN stiffness increased by adding more calcium ionic crosslinker and stress relaxation enhanced by lowering the molecular weight of the alginate. Thus, matrix stiffnesses from 1kPa to 2.5kPa and stress relaxation times from ∼100 s to ∼2,000 s were achieved, values all within physiologically relevant ranges(10, 29).

Here we find that monocytes migrate through the collagen-rich matrices using a novel amoeboid mode of migration. Cells typically migrate in three dimensions utilizing mesenchymal or amoeboid modes of migration. Mesenchymal modes of migration typically involve spread cell morphologies, protrusions, and involve secretion of proteases combined with contractile forces at the protrusive front to generate a path for migration in confining matrices. Further, mesenchymal migration relies on strong cell-matrix adhesion while amoeboid migration uses weak adhesions. 3D migration of monocytes involved rounded or ellipsoidal morphologies and did not rely on strong adhesions, suggesting the migration mode to be more amoeboid in nature. However, the lack of co-localization of myosin and actin differed from known modes of amoeboid migration.

Our 3D migration studies in highly confining hydrogels tested the ability of monocytes to generate migration paths, an ability which is often not assessed in other assays. Cells were embedded in three-dimensional IPN matrices that are nanoporous, therefore requiring cells to generate micron-sized openings to migrate through. Thus, to migrate, encapsulated cells must either degrade the matrix or use mechanical force to deform or remodel the matrix. This contrasts many recent studies that monitor cell migration through microchannels in microfluidic devices, geometries where the migration path is pre-existing, or type-1 collagen matrices with sufficiently large pore sizes(13, 17, 19, 42, 46–56). It is likely that monocytes in vivo do not always have pre-existing paths to migrate through, as the stromal matrix is continuously being remodeled and maintained by fibroblasts, and that some dense stromal matrices they encounter exhibit pore sizes on the nanometer scale, suggesting the physiological relevance of migration path generation. As the IPNs used here contain a nanoporous alginate mesh, and as alginate is not susceptible to degradation by mammalian proteases, the robust 3D migration of the monocytes in the alginate-collagen IPNs occurs independent of proteases. This indicates that monocytes use mechanical forces to mechanically deform and remodel the matrix and generate a migration path, similar to recent observations with cancer cells and mesenchymal stem cells(57, 58). Cancer cells were found to utilize invadopodia protrusions to generate a migration path while mesenchymal stem cells utilize a nuclear piston. Monocytes instead generate substantial protrusive forces using actin polymerization via a Cdc42-WASp signaling axis at the leading edge. These protrusive forces could act to rupture the matrix and create a migration path via divergence of the stresses due to curvature or compression failure in the matrix. Interestingly, this finding of elevated WASp across the leading edge contrasts with the findings of a recent paper, which reported the role of WASp in promoting the generation of small actin patches on the sides of dendritic cells that protrude to help create space for migration in a confining interface(37). Overall, our findings reveal a mechanism of migration path generation in confining matrices that enables monocyte migration.

Furthermore, the finding that monocytes possess the ability to migrate independent of adhesions is consistent with migration studies of other immune cells, though the specific nature of the cell-matrix interactions occurring during monocyte migration may be unique. Previous studies demonstrated that T cells, macrophages, and neutrophils can migrate without adhesions in microfluidic channels with serrated edges or when confined between two macroscopic gels or between glass slides (16, 46, 59, 60). Integrin-independent migration of dendritic cells through collagen and fibrin matrix has also been reported(15). Furthermore, dendritic cells migrate in an adhesion-independent manner using protrusive actin flowing at the leading edge, similar to monocytes, though this has only been shown in microporous gels (15). Here, we show that monocytes migrate with similar characteristics in non-adhesive (alginate only), viscoelastic matrices as they do in collagen-rich matrices. However, our data also show that myosin is not exclusively localized at the rear of the cell as observed during amoeboid migration but is instead distributed throughout the cytoplasm. T cells can also migrate in an amoeboid, adhesion-independent manner using topographical features of the matrix, and the effective friction these features generate on the sides of the cell, to propel cells forward(46). Notably, this mode of migration would result in matrix traction strains parallel to the cell body along the sides, a feature that was not consistently found with monocytes. Monocytes instead mostly typically push off their trailing edge and their sides. All together, our data suggest that monocytes migration in nanoporous matrices is primarily mediated by protrusive forces generated at the leading edge by polymerization of a dendritic actin network, actin flow to the rear of the cell, and cells pushing off their very back and sides.

Changes in extracellular matrix (ECM) stiffness and viscoelasticity are associated with pathological conditions. In addition, mounting evidence implicates migration and accumulation of monocytes and macrophages in disease progression. However, the potential role of ECM changes on monocyte 3D migration has not been reported. Previous work investigated the role of biochemical cues on monocyte/macrophage migration. Our findings on single cell migration complement previous studies that investigated the impact of viscoelasticity on cell clusters (61, 62). Generally, this study raises the possibility that enhanced ECM stiffness and stress relaxation promote monocyte recruitment and migration. However, monocytes could become less migratory as they differentiate into more adhesive macrophage phenotypes. Differences in migration behavior suggests that different molecular targets will likely need to be considered depending on the whether the therapeutic goal is monocyte-depletion or macrophage-depletion. More broadly, our material system provides a platform to study the role of viscoelasticity on migration of normal leukocytes and diseased leukocytes such as those with Leukocyte Adhesion Deficiency-1. Taken together, our data raises the possibility that ECM stiffness and viscoelasticity could determine immune cell recruitment and ultimately shape the immune response under normal and pathological conditions.

## Materials and Methods

### U937 cell culture reagents, primary human cells, and generation of cell lines

U937 cells were maintained in suspension culture in RPMI media containing 2mM glutamine (Thermo Fisher) supplemented with 10% fetal bovine serum (FBS) (Hyclone), and 1% Penicillin/Streptomycin (Life Technologies). Cells were cultured in a standard humidified incubator at 37 °C in a 5% CO_2_ atmosphere. Cells were maintained at sub-confluency and passaged every 2-3 days. To generate frozen aliquots, cells were pelleted by centrifugation (150g, 5min, room temperature), suspended in 90% FBS and 10% dimethylsulfoxide (DMSO, Tocris Bioscience), and frozen in cell-freezing containers at −80°C overnight before transfer to liquid nitrogen for long-term storage. Primary human monocytes were isolated from Leukopak following manufacturer protocols for the EasyEight separation magnet (EasySep^TM^ human CD14 positive selection kit II, STEMCELL^TM^ technologies). The generation of U937 CRISPR KO cell lines were described previously (63, 64). Briefly, 10-sgRNA-per-gene CRISPR/Cas9 deletion library were synthesized, cloned, and infected into Cas9-expressing U937 cells. Puromycin selection (1 μg ml^-1^) was applied to cells for 5 d. Puromycin was then removed and cells were resuspended in normal growth media (no puromycin). Flow cytometry was used to confirm sgRNA infection as >90% of cells were mCherry-positive. Knockout was confirmed using Western Blots. U937 KO lines were stored in liquid nitrogen.

### Alginate preparation

Low molecular weight (MW) ultra-pure sodium alginate (Provona UP VLVG, NovaMatrix) was used for fast-relaxing substrates, with MW of <75 kDa, according to the manufacturer. Sodium alginate rich in guluronic acid blocks and with a high-MW (FMC Biopolymer, Protanal LF 20/40, High-MW, 280 kDa) was prepared for slow-relaxing substrates. Alginate was dialyzed against deionized water for 3–4 days (MW cutoff of 3,500 Da), treated with activated charcoal, sterile-filtered, lyophilized, and then reconstituted to 3.5 wt% in serum-free Dulbecco’s modified Eagle’s medium (DMEM, Gibco). The use of low/high molecular weight alginate resulted in fast/slow-relaxing IPNs.

### Mechanical characterization of IPNs

IPNs were characterized as previously described(65). Briefly, rheology testing was done with a stress-controlled AR2000EX rheometer (TA instruments). IPNs for rheology testing were deposited directly onto the bottom Peltier plate. A 25 mm flat plate was then slowly lowered to contact the gel, forming a 25 mm disk gel. Mineral oil (Sigma) was applied to the edges of the gel to prevent dehydration. For modulus measurement, a time sweep was performed at 1 rad/s, 37 °C, and 1% strain for 3 h after which the storage and loss moduli had equilibrated. Young’s modulus (E) was calculated, assuming a Poisson’s ratio (v) of 0.5, from the equation:

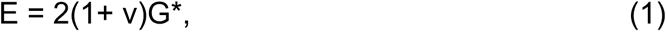

where complex modulus, G*, was calculated from the measured storage (G’) and loss moduli (G’’) using:

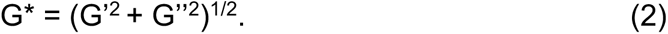

For stress relaxation experiments, the time sweep was followed by applying a constant strain of 5% to the gel, at 37 °C, and the resulting stress was recorded over the course of 3 h.

### Hydrogel formation, cell encapsulation and monocyte differentiation

For each viscoelastic gel, alginate was delivered to a 1.5 mL eppendorf tube (polymers tube) at room temperature. Rat tail collagen I (Corning), was neutralized with 10X DMEM and pH adjusted to 7.4. Neutralized collagen was added to the alginate and carefully mixed 30 times with a pipette, being careful not to generate bubbles. Extra DMEM was added to ensure 4.8 mg/ml - 1.6 mg/ml alginate-collagen final gel concentration. For 3D migration assays, cells were resuspended in growth media containing octadecyl rhodamine B chloride (R18, ThermoFisher, 1:1000 dilution of 10mg/ml stock), centrifuged, and re-suspended in growth media. The concentration of cells was determined using a Vi-Cell Coulter counter (Beckman Coulter) after passing through a 40 μM filter (Fisher Scientific) to obtain single cell suspensions. ∼2×10^6^ cells were encapsulated in each gel condition. Extra DMEM was added such that all substrates had a final concentration of 4.8 mg/mL alginate and 1.6 mg/mL collagen. This was mixed 30 times with a pipette.

Next, different calcium sulfate concentrations were added to a 1 mL Luer lock syringe (Cole-Parmer), to ensure that the initial Young’s modulus is kept constant for fast, and slow-relaxing substrates. The mixture of the polymers was transferred to a separate 1 mL Luer lock syringe (polymers syringe). The calcium sulfate solution was shaken to mix the calcium sulfate evenly, and it was then coupled to the polymers syringe with a female-female Luer lock (Cole-Parmer), taking care not to introduce bubbles or air in the mixture. Finally, the two solutions were rapidly mixed together with 15 pumps on the syringe handles and instantly deposited into a well in an 8-well Lab-Tek dish (Thermo Scientific). We sought to maintain similar collagen fiber architecture when IPN stiffness and viscoelasticity was tuned. The optimal conditions we found to accomplish this was initiation of IPN gelation at 22°C for 1 h before media was added to the wells. The samples were then allowed to gel for an additional 1 h followed by transfer to 37°C incubator to complete gelation. Media was replaced with fresh media for all gels 24 h after encapsulation. For cell differentiation assays, 200 ng/ml Phorbol 12-myristate 13-acetate (Fisher Scientific, PMA) was added to the gel.

### Inhibition studies

Pharmacological inhibitors were added to cell media 10 minutes before time-lapse microscopy experiments. The concentrations used for the inhibitors are: 2 μM Latrunculin A (Tocris Bioscience, actin polymerization inhibitor), 100 μM CK-666 (Sigma, Arp 2/3 actin polymerization inhibitor), 100 μM Y-27632 (Sigma, ROCK inhibitor), 50 μM MLCK (Sigma, myosin inhibitor), 2 μM DDR1 (Fisher Scientific, discoidin domain receptor 1 inhibitor), 40 μM EIPA (Sigma, Na+/H+ exchange inhibitor), 20 μM ML141 (Tocris Bioscience, Cdc42 GTPase inibitor), 25 μM GSK205 (Calbiochem, TRPV4 antagonist), and 10 μM fasudil (Stemcell, ROCK-dependent contractility inhibitor). Time lapse images were acquired every 10 minutes for 24 hours.

### Live/dead studies

We determined live and dead cells using the commonly used LIVE/DEAD viability/Cytotoxicity kit (Fisher Scientific). We incubated U937 with 2 μM of calcein-AM dye and 4μM of ethidium homodimer-1 for 45 minutes and then immediately imaged the cells. Live cells exhibit green fluorescence indicating esterase activity while dead cells exhibit red fluorescence due to loss of plasma membrane integrity.

### Immunofluorescence for fixed cells

Cells were embedded in matrix for 24 hours. Media was then removed from the matrix and replaced with low-melting-temperature agarose to prevent matrix from floating in subsequent steps. Matrix was then washed with serum-free DMEM and then fixed with 4% paraformaldehyde in serum-free DMEM, at room temperature, for 20 min. This was followed by three washes with Phosphate Buffered Saline (PBS) for 10 min each time. IPN hydrogels were then embedded in OCT compound (optimal cutting temperature compound; Fisher Scientific) and cryo-sectioned. Cells were then permeabilized with a permeabilizing solution for 15 min and washed twice with PBS for 5 min each time. Blocking solution was added to minimize non-specific staining. After this, primary antibodies were added overnight at room temperature and subsequently washed twice with PBS. Secondary antibodies, DAPI and phalloidin, were added for 1.5 h at room temperature followed by two PBS washes. ProLong Gold antifade reagent (Life Technologies) was added just before imaging to minimize photobleaching. Images were acquired with a Leica 63X objective.

### Western blot experiments

For sample preparation, cells were incubated in RIPA Lysis and Extraction Buffer (Thermo Scientific) containing Halt^TM^ Protease and Phosphatase Inhibitor Cocktail (Thermo Scientific) on ice for 20 min. The samples were centrifuged at 4 ° C for 15 min at 14,000 g, and the supernatant was collected. The protein concentration of the lysate was quantified using the Pierce^TM^ BCA Protein Assay Kit (Thermo Scientific). 10 ug of protein was loaded onto NuPAGE^TM^ Bis-Tris protein gel (Invitrogen) for β_2_-integrin blotting or NuPAGE^TM^ Tris-Acetate protein gel (Invitrogen) for talin-1 blotting. The lysates were separated by SDS-PAGE and transferred to Immobilon-P PVDF Membrane (Milipore). The membranes were incubated in No-Stain^TM^ Protein Labeling Reagent (Invitrogen) for 10 min for total protein labeling and washed 3 times with water. The membranes were then blocked in blocking solution (0.1% non-fat dry milk in tris-buffered saline with Tween-20, TBST) at room temperature for 1 h. Primary antibodies (β_2_-integrin, Proteintech, 10554-1-AP; tailn-1, Cell Signaling Technology, 4021) were diluted 1:2,000 in blocking solution, and the membranes were incubated with the primary antibodies overnight at 4 ° C. The membraned were then washed 3 times with TBST for 5 min each and incubated in HRP-conjugated goat anti-rabbit antibody (Abcam, ab6721, 1:100,000) in blocking solution at room temperature for 1 h. The membranes were then washed with TBST, developed with SuperSignal^TM^ West Femto Maximum Sensitivity Substrate (Thermo Scientific), and imaged using the iBright^TM^ CL1500 imaging system (Invitrogen). Band intensities were quantified using ImageJ.

### Confocal microscopy

Microscope imaging was done with a laser scanning confocal microscope (Leica SP8) and a spinning disk confocal microscope (Nikon Eclipse Ti2). Both microscopes were fitted with temperature/incubator control, suitable for live imaging (37 °C, 5% CO_2_). In live-cell time-lapse migration imaging, R18 membrane labeled cells were tracked with a Leica 20X NA = 0.75 air objective for 20 hours. 50 μm stack images were acquired every 10 minutes and imaging parameters were adjusted to minimize photobleaching and avoid cell death. For actin flow experiments, U937 monocytes were incubated with 1:2,000 dilution factor SPY650-FastAct (Cytoskeleton, Inc., SC505, F-actin stain) for 8 hours, and 20 μm image stacks were acquired every 60 seconds with a Nikon 40X NA = 1.15 oil objective for 30 minutes. For WASp-GFP experiments, 50 μm image z-stacks were acquired every 2 minutes with a Leica 25X NA = 0.95 water objective for 40 minutes. Heatmaps were generated by taking the maximum intensity z-projection, applying a smoothing algorithm, and coloring cells using the 16-color lookup table within ImageJ.

### Cell volume measurements

U937 monocytes were incubated with 1:2,000 dilution factor SPY650-FastAct (Cytoskeleton, Inc., F-actin stain) for 12 hours. 30 μM image stacks were acquired every 60 seconds with a Nikon 40X NA = 1.15 oil objective for 30 minutes. Individual cells that transition from stationary to migratory were identified and manually segmented within Imaris to obtain volume measurements.

### Confocal reflectance microscopy for collagen fiber characterization

Alginate-collagen matrices in 8-well Labtek chamber were mounted on a laser scanning confocal microscope (Leica SP8) equipped with a 25X NA = 0.95 water-matched objective. A single slice of the sample was excited at 488 nm and reflected light was collected using the “Reflectance” setting on the microscope. Several, randomly selected sections of the matrix were imaged. Collagen fibril length and width were analyzed using the free and publicly available CT-FIRE software (http://loci.wisc.edu/ software/ctfire)(66–68).

### Imaris cell tracking algorithm

For migration studies, the centroids of R18-labeled cells were tracked using the spots detection functionality in Imaris (Bitplane). Poorly segmented cells and cell debris were excluded from the analysis and drift correction was implemented where appropriate. A custom MATLAB script was used to reconstruct cell migration trajectory.

### Measuring cell-induced ECM deformations

To measure ECM mechanical deformations induced by cell forces, IPNs were seeded with fluorescent beads following established protocols(25, 65). For 3D deformation field, fast-iterative digital volume correlation(69) with a subset size of 32×32×32 voxels, and spacing of 8 voxels (2.4 μm), was employed. For 2D deformations, fast-iterative digital image correlation(70) was used on confocal images of fluorescent beads with a subset size of 32×32 pixels, and spacing of 8×8 pixels (2.4 um). To average 2D deformation field across various cells, we measured the direction of velocities of the cells, and rotated the deformation field accordingly, with each cell’s velocity directed along the “x-axis”. This enabled us to average the ECM deformations around the cell with respect to the direction of cell movement.

### WASp-GFP reporter generation

To generate a U937 cell line stably expressing a WASp protein fused to monomeric enhanced green fluorescent protein (mEGFP), a lentiviral vector encoding the mEGFP-WASp fusion gene under the control of the spleen focus-forming virus (SFFV) constitutive promoter was constructed. The full coding sequence of human WASp was retrieved from the GRCh38 human genome assembly using Benchling. The mEGFP sequence was obtained from the mEGFP-C1 vector (a gift from Michael Davidson; Addgene plasmid # 54759). A flexible linker consisting of two repeats of the amino acid sequence GGGGS [(GGGGS)₂] was inserted between the C-terminus of mEGFP and the N-terminus of WASp. This linker was incorporated to ensure proper folding and functionality of the fusion protein while preserving the integrity of the VCA domain of WASp, which interacts with the ARP2/3 complex. The mEGFP-(GGGGS)₂-WASp fusion gene was synthesized and cloned into a lentiviral vector backbone by Twist Bioscience (San Francisco, CA, USA).

Lentiviral particles were produced in HEK293T cells using second-generation packaging plasmids as previously described. Briefly, the transfer plasmid (mEGFP-WASp), packaging plasmid psPax2 (a gift from Didier Trono; Addgene plasmid #12260, and envelope plasmid pMD2.G (a gift from Didier Trono; Addgene plasmid #12259 were mixed at a molar ratio of 2:1:1. The DNA mixture was combined with linear polyethyleneimine (PEI, 25 kDa, Polysciences, Inc.) at a PEI:DNA mass ratio of 2:1 and incubated for 20 minutes at room temperature to form polyplexes. The PEI-DNA complexes were added to HEK293T cells at approximately 50% confluence in Opti-MEM reduced-serum medium (Gibco, Thermo Fisher Scientific). After a 4-hour incubation, the medium was replaced with complete Dulbecco’s Modified Eagle Medium (DMEM) supplemented with 10% FBS and 1% Pen-Strep. The cells were cultured for an additional 48 hours to allow virus production. The virus-containing supernatant was collected, filtered through a 0.45 μm polyethersulfone (PES) filter, and concentrated approximately 50-fold using the Lenti-X Concentrator Kit (Takara Bio USA) according to the manufacturer’s protocol. U937 cells were seeded at 400,000 cells/mL in complete RPMI 1640 medium supplemented with 10% FBS and 1% Pen-Strep. Thawed viral stocks were added to the cells, and spinoculation was performed at 1,000 × g for 90 minutes at 32 °C in the presence of polybrene (5 μg/mL, Sigma-Aldrich). Following spinoculation, the cells were incubated at 37 °C with 5% CO₂ for 24 hours. The cells were then washed twice with RPMI 1640 medium to remove residual viral particles and cultured in fresh complete RPMI 1640 medium. To maintain high levels of expression and prevent potential silencing or loss of the construct, live-cell imaging was performed within two passages post-transduction.

### Statistics and reproducibility

All measurements were performed on 2-3 biological replicates from separate experiments, with at least 2 biological replicates for data with conditions that are compared statistically for significant differences. Exact sample size and exact statistical test performed for each experiment is indicated in appropriate figure legends. Statistical analyses were performed using GraphPad Prism (La Jolla, California). For all violin plots, broken lines are median values. For statistical comparisons, we treated each cell as an independent entity and averaged across all experiments and tested for statistics. For scatter plots, solid lines are median values. p-values reported were corrected for multiple comparisons, where appropriate.

## Supporting information

Adebowale Monocyte SI_December 2024_FINAL_bioRxiv

## Data availability

All data relevant to this manuscript will be made available as a supplementary text file or deposited in a public repository.

## Acknowledgments

We acknowledge members of the Chaudhuri lab for helpful discussions and Marc Levenston (Stanford University) for use of mechanical testing equipment. We thank Vivek B. Shenoy (University of Pennsylvania) for insightful discussions. We also acknowledge the Stanford Cell Sciences Imaging Facility for Imaris software access and for technical assistance with Imaris. We thank Yael Rosenberg-Hasson for performing the flow cytometry experiments at Stanford’s Human Immune Monitoring Center (HIMC). K.A. acknowledges financial support from the Stanford ChEM-H Chemistry/Biology Interface Predoctoral Training Program and the National Institute of General Medical Sciences of the National Institutes of Health under Award Number T32GM120007, and a National Science Foundation Graduate Student fellowship. This work was supported by National Institutes of Health National Cancer Institute Grant (R37 CA214136) and NSF CAREER award (CMMI 1846367) for O.C.

## Supporting Information for

**Fig. S1.**
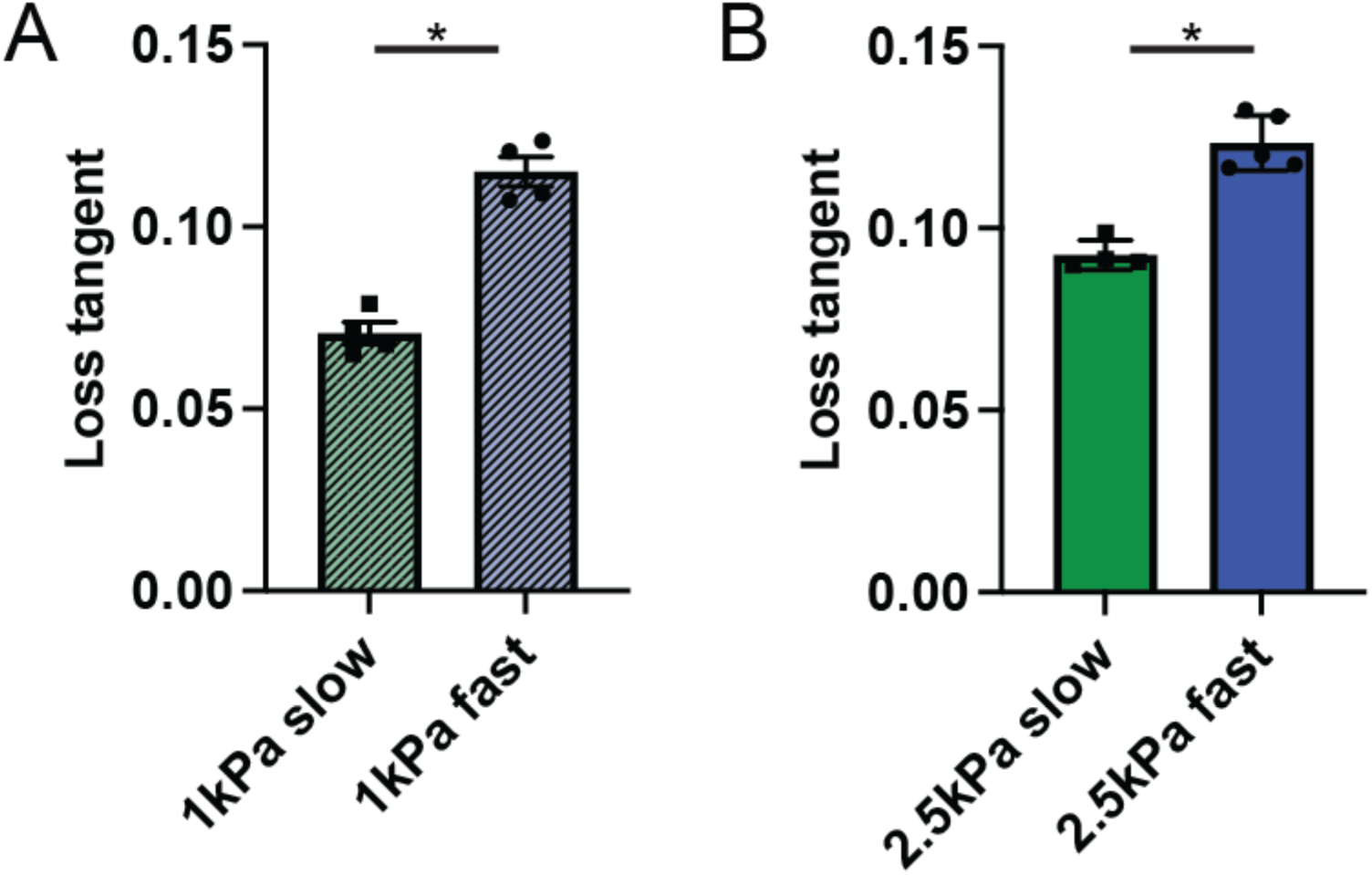
Loss tangent measurements of the different IPN formulations for slow and fast matrices. (A) N = 4 biological replicates for each condition, Kolmogorov-Smirnov test: * p = 0.0286. (B) N ≥ 4 biological replicates for each condition, Kolmogorov-Smirnov test: * p = 0.0159.

**Fig. S2.**
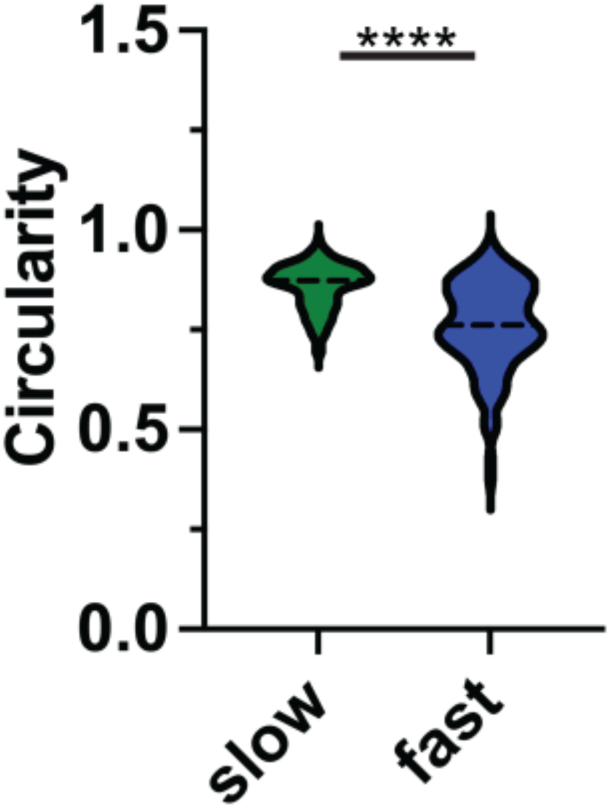
Circularity for U937 human monocytes embedded in slow and fast IPN matrices. n = 132 for each condition, N = 2 biological replicates. Kolmogorov-Smirnov test: **** p < 0.0001.

**Fig. S3.**
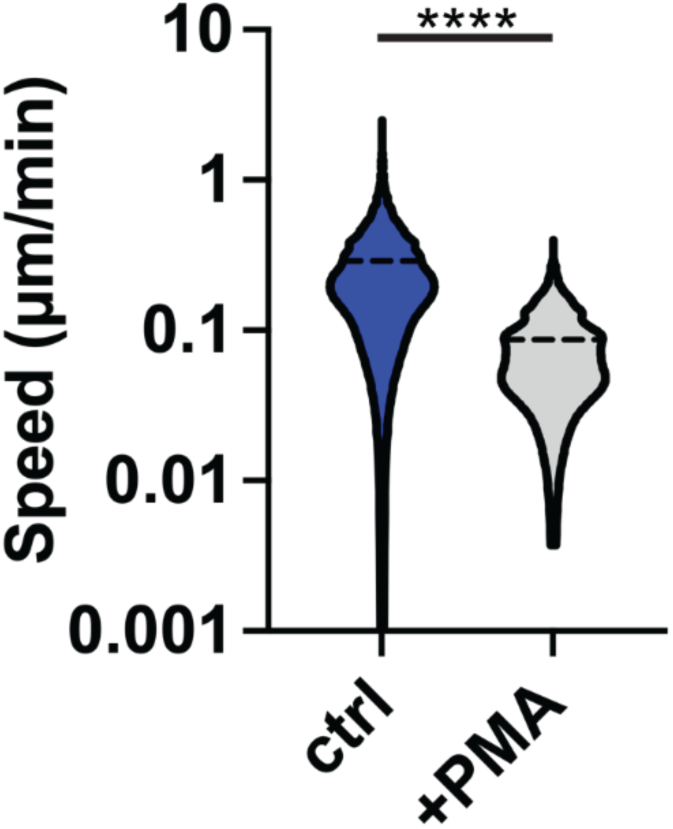
U937 migration decreases upon addition of Phorbol 12-myristate 13-acetate (PMA) to U937 cells embedded in fast relaxing IPN matrix. n > 790, N = 2 biological replicates. Kolmogorov-Smirnov test: **** p < 0.0001.

**Fig. S4.**
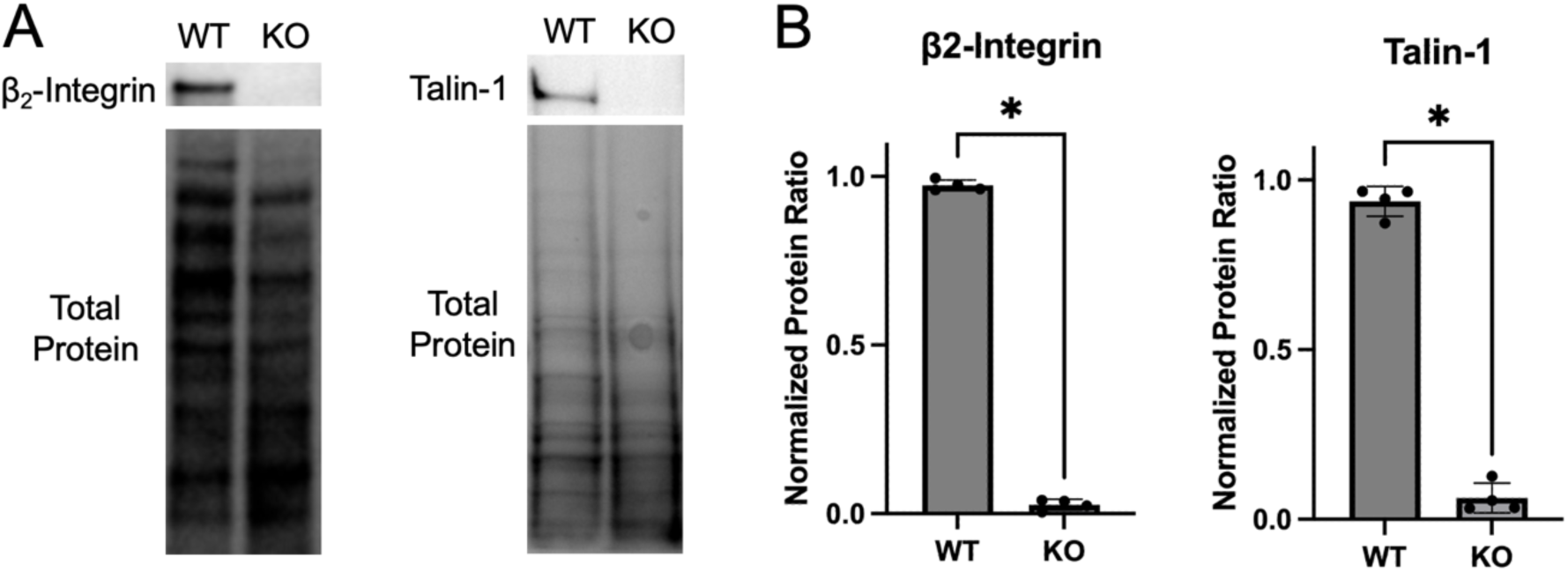
Western blot analysis confirming CRISPR knockouts (KO) of β2-integrin and talin-1 in U937 cells. (A) Western blot showing wild type (WT) and KO cells. (B) Quantification of western blot results across 4 experiments and 2 biological replicates for each condition. Kolmogorov-Smirnov test: * p < 0.05.

**Fig. S5.**
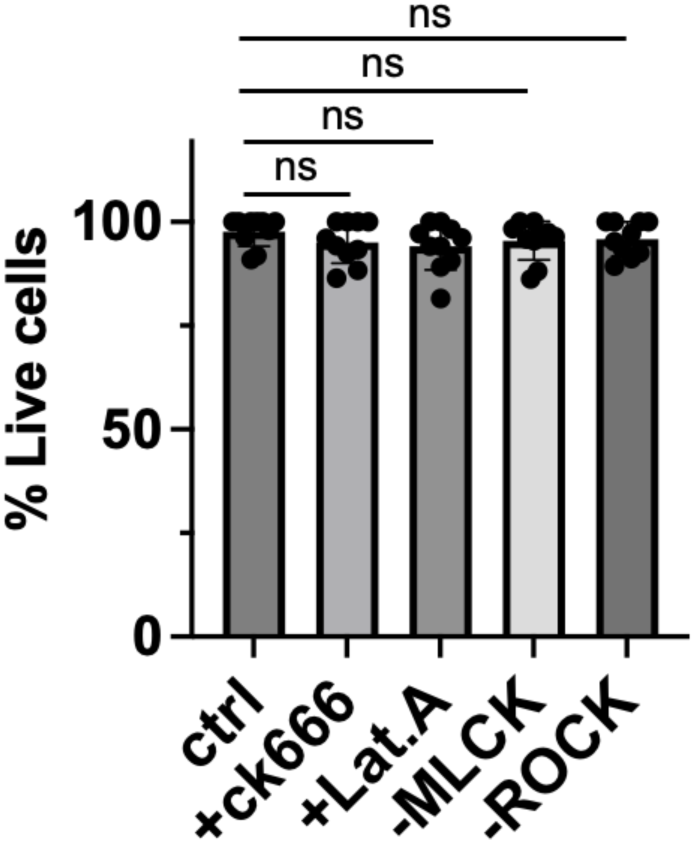
Effect of pharmacological inhibition on cell viability. (A,B) n > 218 for each condition; N = 2 biological replicates. Kruskal-Wallis test with Dunn’s multiple comparisons: ns p > 0.9999.

**Fig. S6.**
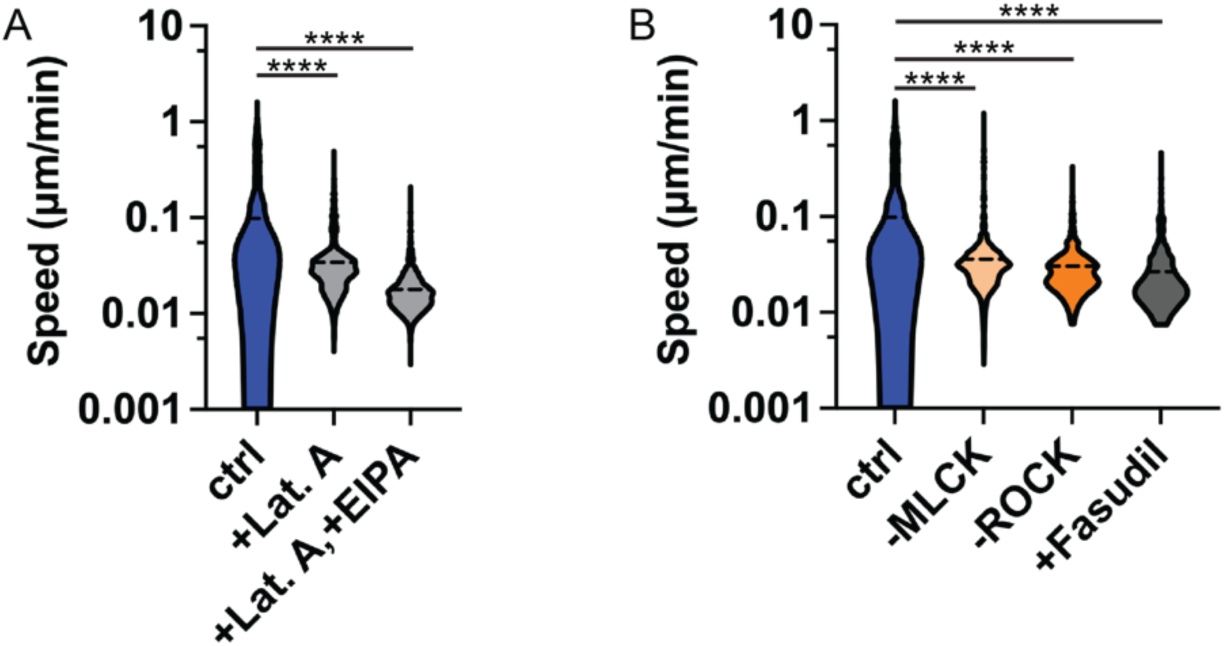
Effect of actin, myosin, and contractility on speed of migration for primary human monocytes embedded in fast IPN matrices. (A,B) n > 557 for each condition; N > 2 biological replicates. Kruskal-Wallis test with Dunn’s multiple comparisons: ns p > 0.9999, **** p < 0.0001.

**Fig. S7.**
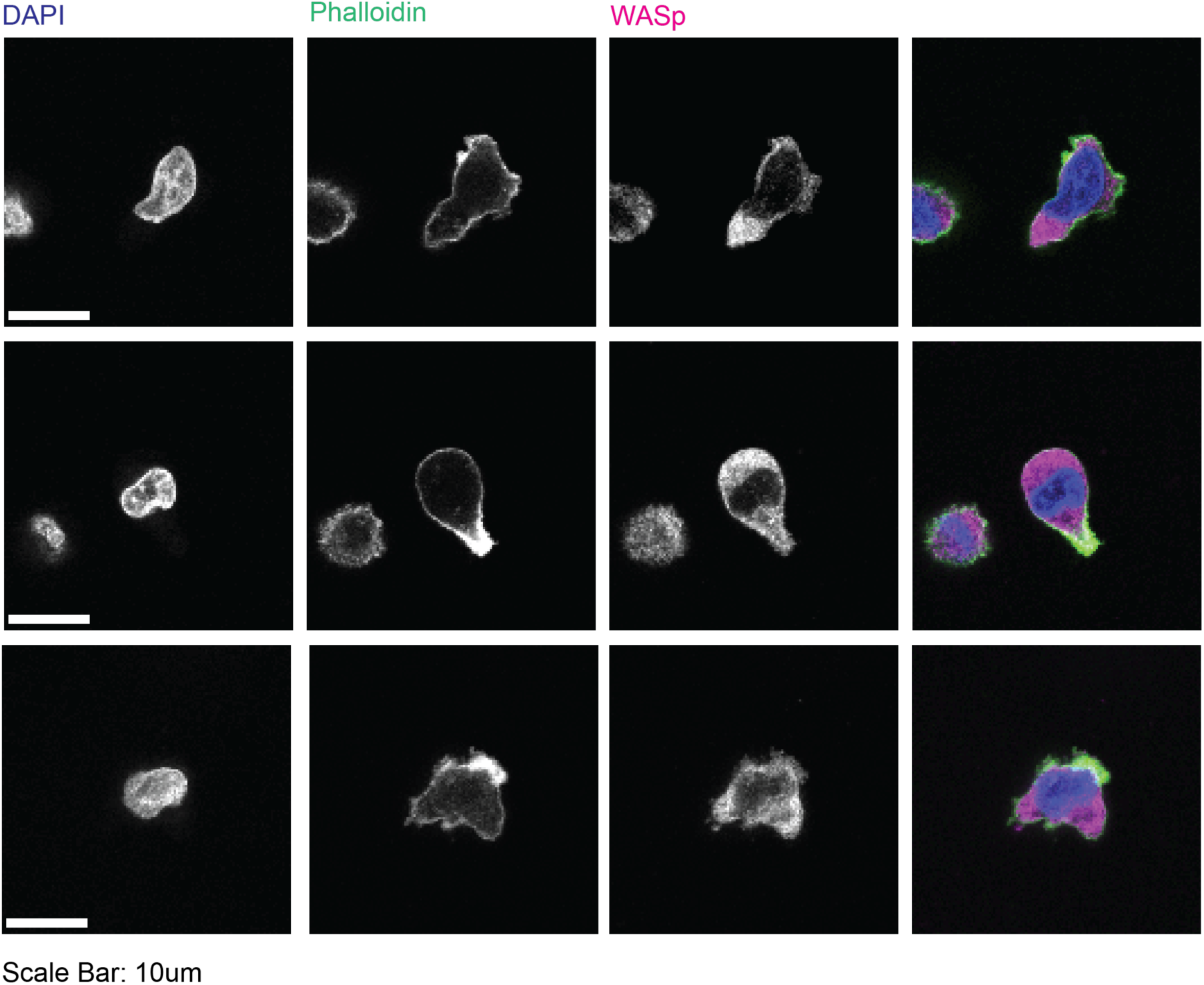
(A,B) Immunofluorescence images of representative cells staining nucleation promotion factor WASp, along with DAPI (nucleus) and phalloidin (actin filaments). Scale bar: 10 μm. These data are from U937 cells.

**Fig. S8.**
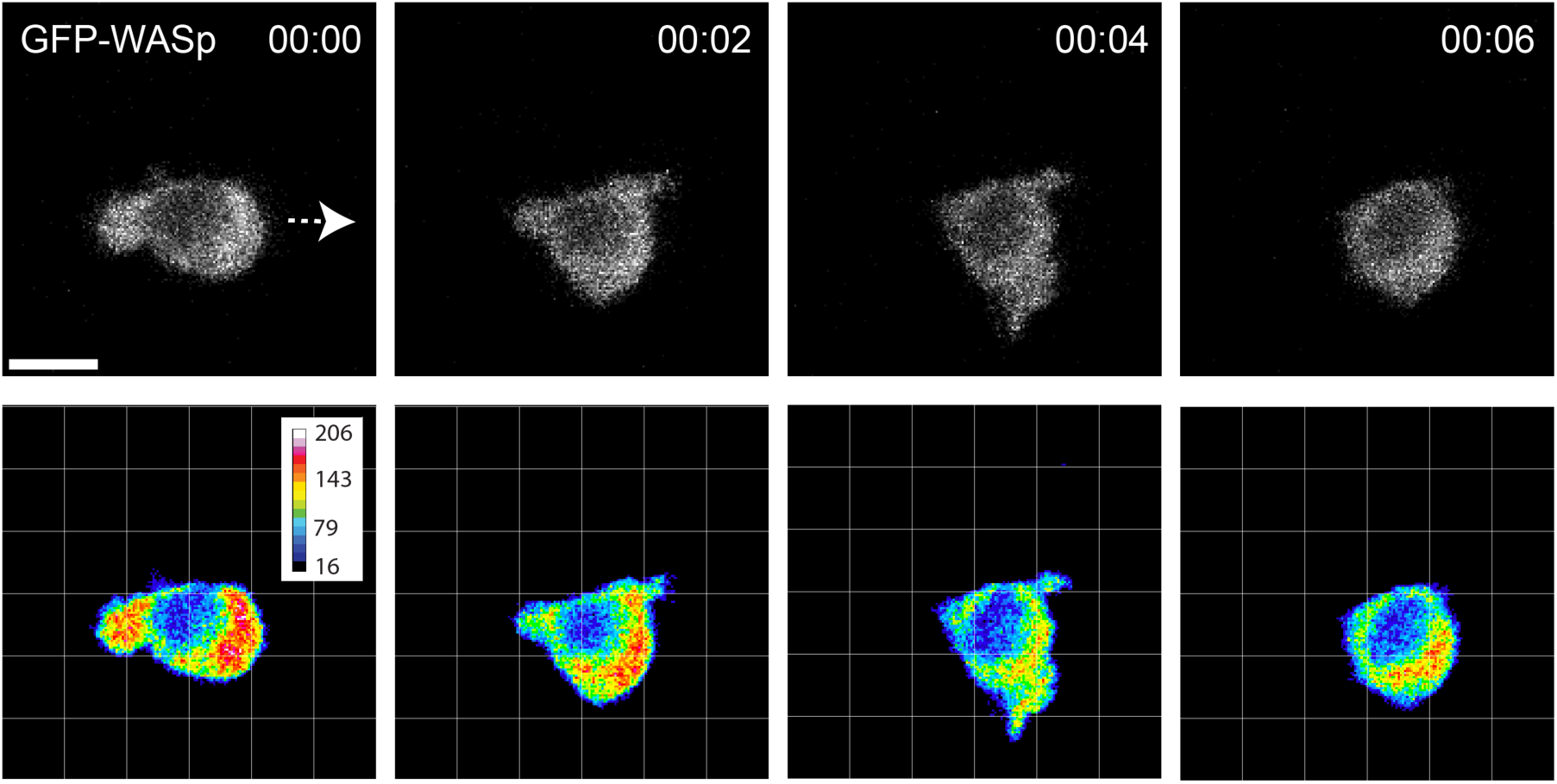
Live imaging of fluorescently labeled WASp at the front of the cell. White arrow indicates migration direction. Scale bar: 10 μm. These data are from U937 cells.

**Fig. S9.**
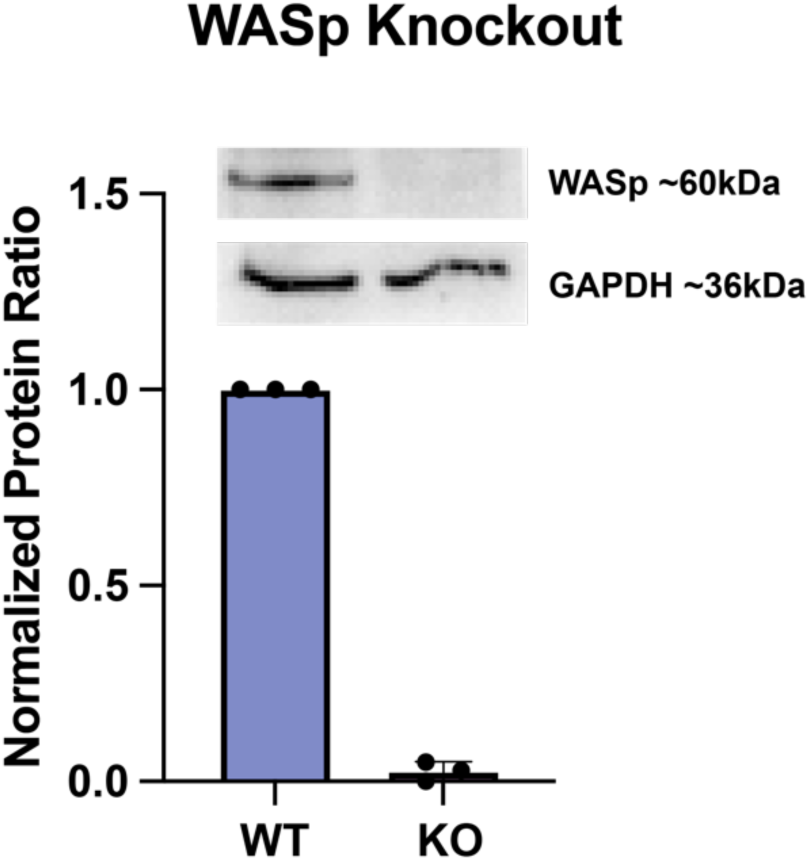
Western blot analysis confirming CRISPR knockouts (KO) of WASp and quantification in U937 cells. N = 3 biological replicates.

**Fig. S10.**
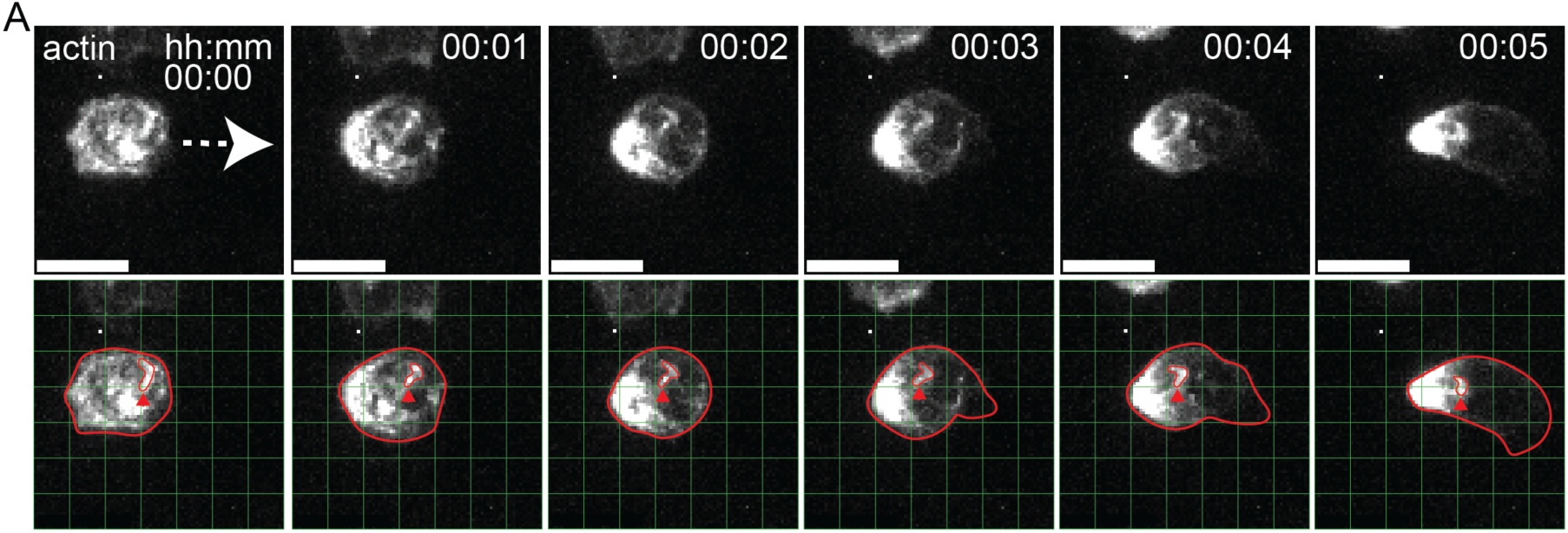
Maximum intensity z-projections of actin in a migrating U937 monocyte. Time interval between succussive frames is 60 seconds. For top row, white arrow indicates direction of migration with actin puncta at cell rear. For bottom row of images, cell and actin spot are outlined in red, and a green grid superimposed to serve as a spatial reference. Red arrow shows retrograde flow of actin puncta.

