## Supplementary material for "Monocytes use protrusive forces to generate migration paths in viscoelastic collagen-based extracellular matrices": Adebowale Monocyte SI_December 2024_FINAL_bioRxiv: Adebowale Monocyte SI_December 2024_FINAL_bioRxiv.pdf

**This PDF file includes:**

Figures S1 to S10

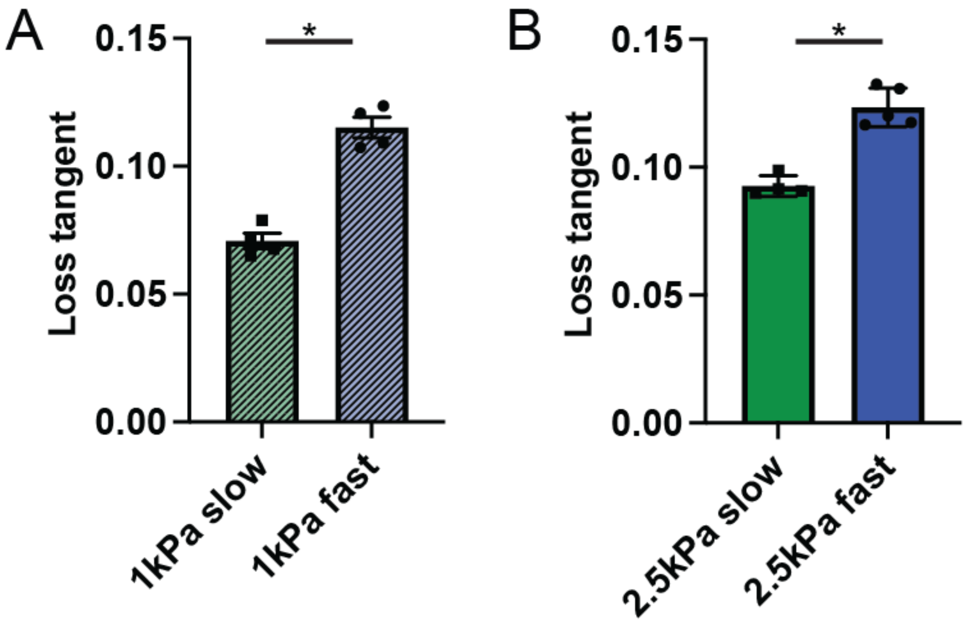


**Fig. S1** Loss tangent measurements of the different IPN formulations for slow and fast matrices. (A) N = 4 biological replicates for each condition, Kolmogorov-Smirnov test: * p = 0.0286. (B) N $\geq$ 4 biological replicates for each condition, Kolmogorov-Smirnov test: * p = 0.0159.

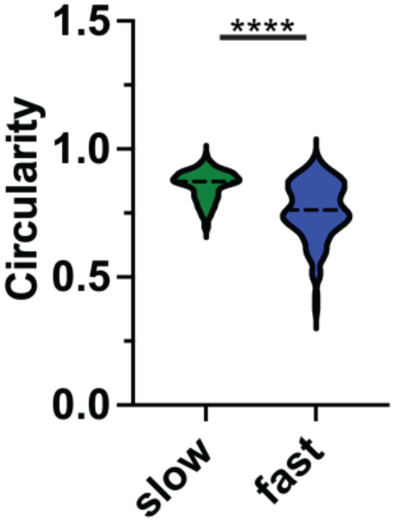


**Fig. S2** Circularity for U937 human monocytes embedded in slow and fast IPN matrices. n = 132 for each condition, N = 2 biological replicates. Kolmogorov-Smirnov test: **** $p < 0.0001$.

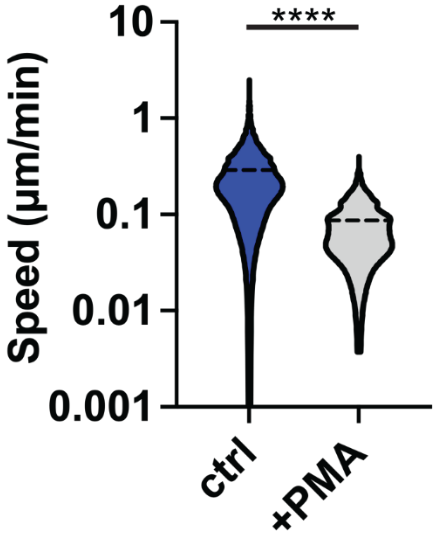


**Fig. S3** U937 migration decreases upon addition of Phorbol 12-myristate 13-acetate (PMA) to U937 cells embedded in fast relaxing IPN matrix. n > 790, N = 2 biological replicates. Kolmogorov-Smirnov test: **** $p < 0.0001$.

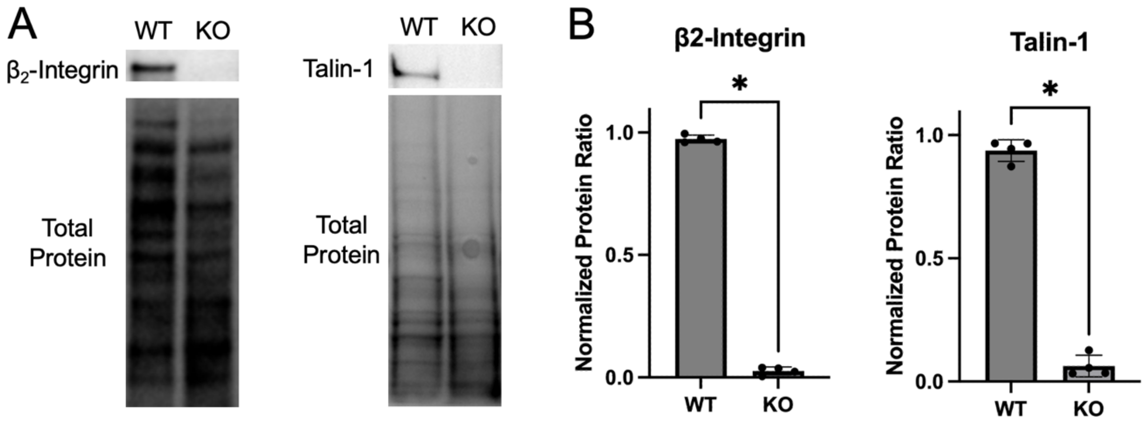


**Fig. S4** Western blot analysis confirming CRISPR knockouts (KO) of $\beta_2$-integrin and talin-1 in U937 cells. (A) Western blot showing wild type (WT) and KO cells. (B) Quantification of western blot results across 4 experiments and 2 biological replicates for each condition. Kolmogorov-Smirnov test: * $p < 0.05$.

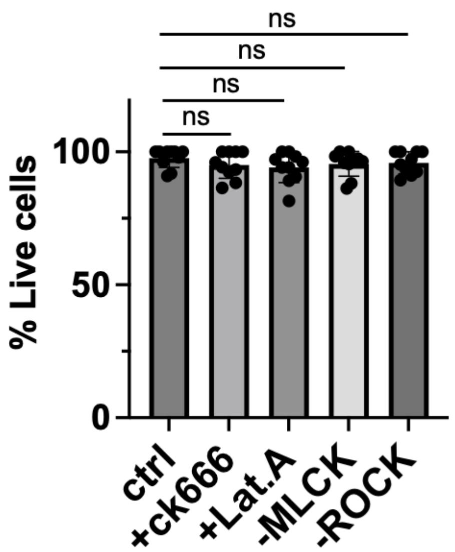


**Fig. S5** Effect of pharmacological inhibition on cell viability. (A,B) n > 218 for each condition; N = 2 biological replicates. Kruskal-Wallis test with Dunn’s multiple comparisons: ns p > 0.9999.

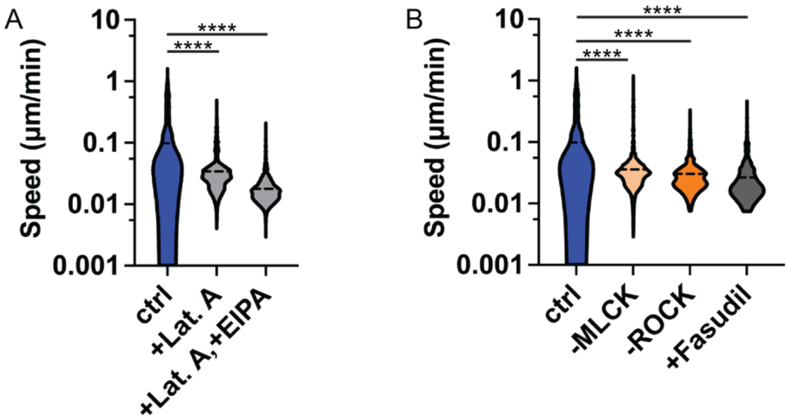


**Fig. S6** Effect of actin, myosin, and contractility on speed of migration for primary human monocytes embedded in fast IPN matrices. (A,B) n > 557 for each condition; N > 2 biological replicates. Kruskal-Wallis test with Dunn's multiple comparisons: ns p > 0.9999, **** p < 0.0001.

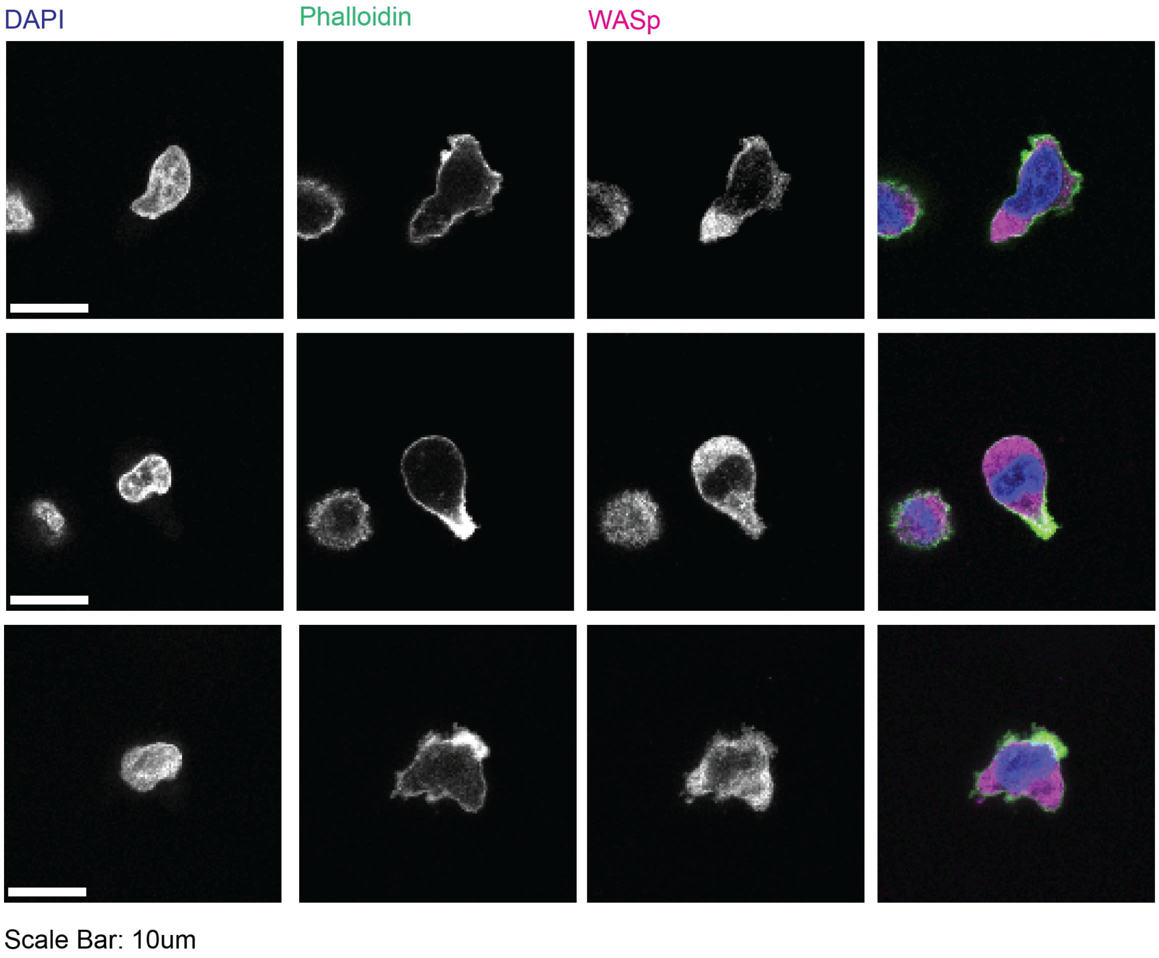


**Fig. S7** (A,B) Immunofluorescence images of representative cells staining nucleation promotion factor WASp, along with DAPI (nucleus) and phalloidin (actin filaments). Scale bar: 10 μm. These data are from U937 cells.

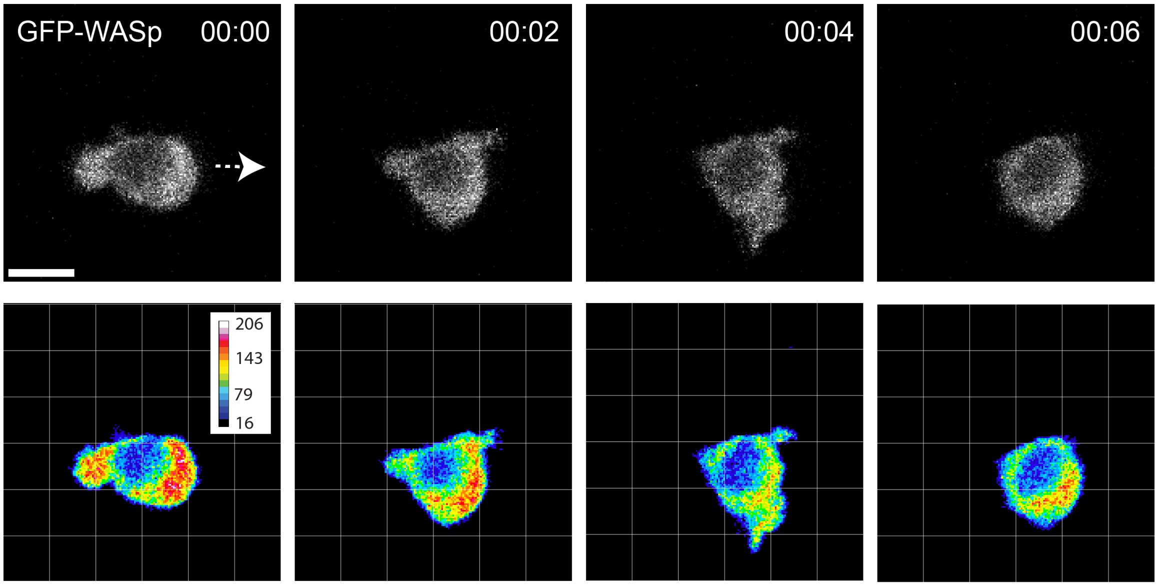


**Fig. S8** Live imaging of fluorescently labeled WASp at the front of the cell. White arrow indicates migration direction. Scale bar: 10 μm. These data are from U937 cells.

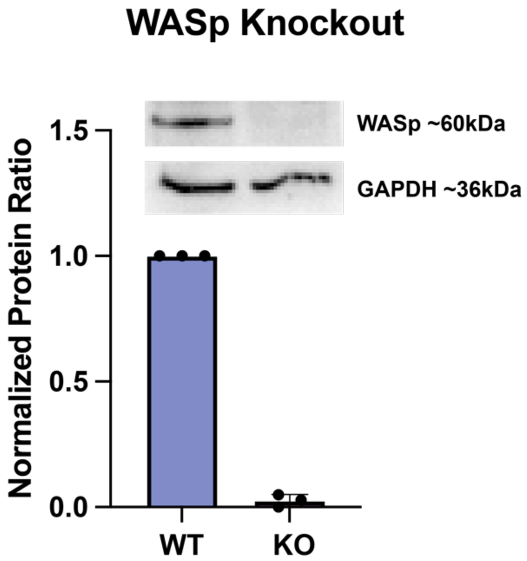


**Fig. S9** Western blot analysis confirming CRISPR knockouts (KO) of WASp and quantification in U937 cells. N = 3 biological replicates.

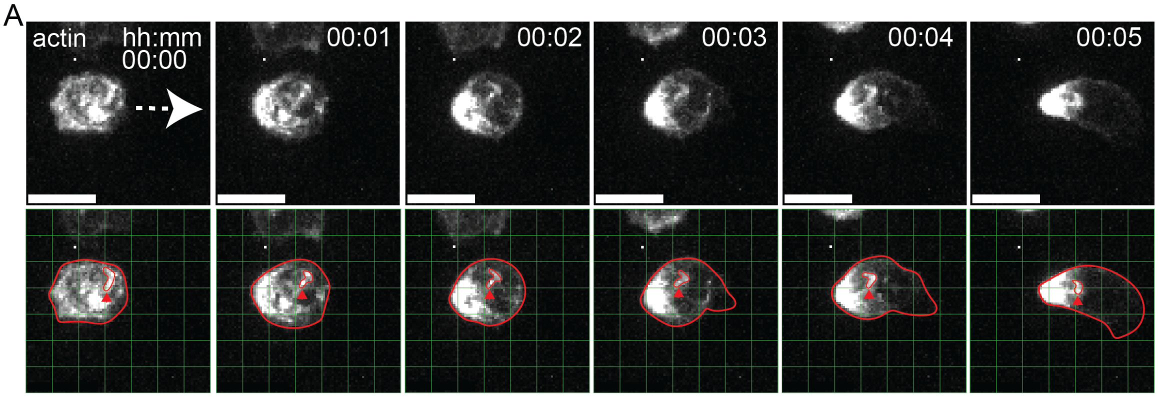


**Fig. S10** Maximum intensity z-projections of actin in a migrating U937 monocyte. Time interval between succussive frames is 60 seconds. For top row, white arrow indicates direction of migration with actin puncta at cell rear. For bottom row of images, cell and actin spot are outlined in red, and a green grid superimposed to serve as a spatial reference. Red arrow shows retrograde flow of actin puncta.
